# Cell-Cell Communication and Gene Regulation of Stem Cell-Parenchymal Cell Fusion

**DOI:** 10.1101/2025.02.11.637745

**Authors:** Fateme Nazaryabrbekoh, JoAnne Huang, Syeda S. Shoaib, Xun Tang, Joohyun Kim, Brenda M. Ogle, Jangwook P. Jung

## Abstract

Cell fusion, a natural process occurring between similar or dissimilar cell types, often confers new or enhanced functionality, yet its impact on cellular communication and gene regulation remains poorly understood. Here, we used recent analytical frameworks to investigate accidental cell fusion between murine cardiomyocytes (mHL1) and murine mesenchymal stromal/stem cells (mMSC) leveraging previously published single-cell RNA sequencing data. After fusion, we observed a biased distribution of gene expression and acquired phenotypes in fused hybrids. Trajectory inference showed that hybrids with a more mMSC-like transcriptome diverged more significantly from parental cells relative to hybrids with a more mHL1-like transcriptome. We also observed dynamic changes in cell-cell communication, with early (Day 1) downregulation of Wnt signaling and Melanogenesis evolving into the upregulation of pathways like Endocrine resistance and Focal adhesion by Day 3. Notably, ECM (extracellular matrix)-receptor interactions were largely consistent whether annotation or unsupervised Clustree methods were used. Furthermore, our analysis indicated the emergence of various cancer-associated signaling mechanisms. Our findings highlight the remarkable plasticity of cellular identity following fusion and lay the groundwork for future research into the precise molecular mechanisms driving these transformations and the potential of cell fusion for generating novel cell types.

## Introduction

Cell-cell fusion is a biological process where two cells merge their membranes, forming a single hybrid cell^1^. This fundamental event plays a crucial role in development, reproduction, and tissue repair across various organisms^2^. Heterotypic fusion, the merging of cells with distinct origins, presents a complex interplay of benefits and risks. Essential for processes such as fertilization, it can also disrupt cellular homeostasis. The fusion of distinct cellular environments can have profound effects on nuclear function, including competition for gene regulation, chromosomal reorganization, and even nuclear ejection^3-5^. Mesenchymal stromal cells (MSCs) exhibit a remarkable capacity to fuse with diverse cell types within various organs, including the brain, liver, and heart^6-13^. This phenomenon of cell fusion plays a pivotal role in development and regeneration, while also contributing to cancer progression. Notably, MSC fusion has demonstrated beneficial effects in certain contexts, such as promoting liver regeneration through fusion with hepatocytes^6,14,15^ and contributing to muscle regeneration through fusion with myocytes^16,17^. However, the consequences of MSC fusion in many other contexts remain uncertain. A growing body of evidence links heterotypic fusion involving MSCs to the development of cancer and metastasis, raising significant concerns about the safety of MSC-based therapies^18-23^. Understanding the underlying mechanisms of cell fusion is crucial to distinguish between beneficial and detrimental outcomes.

This study revisits previously published data on the fusion of murine HL-1 cardiomyocytes (mHL1) and murine MSCs (mMSC)^23^. While the original study utilized bulk RNA-seq analysis methods, we now leverage more recent single-cell RNA-sequencing (scRNA-seq) analysis tools to gain deeper insights into cell-cell communication and the underlying gene networks orchestrating this specific fusion process. By applying contemporary scRNA-seq analysis tools, we aim to extend beyond the previous limitations of principal component analysis (PCA), hierarchical clustering (HC), and Gene Ontology (GO) analyses^23^. **Figure 1** provides a visual overview of our study examining the outcomes of accidental mMSC and mHL1 cell fusion, utilizing scRNA-seq analysis at days 1 and 3 post-fusion. By integrating both annotated and unsupervised clustering approaches, we delineated distinct cell populations arising from these fusion events. Notably, trajectory inference (TI) analysis revealed a temporal evolution in the identity of the fusion products. To elucidate the transcriptional mechanisms driving these changes, we further investigated alterations in both intercellular communication pathways (including Secreted Signaling, ECM-Receptor Engagement, and Cell-Cell Contact) and intracellular communication networks (gene regulatory networks). We then contextualize these new findings within our previous work, offering a more in-depth understanding of cellular communication dynamics across different stages of mHL1-mMSC fusion.

**Figure 1.**
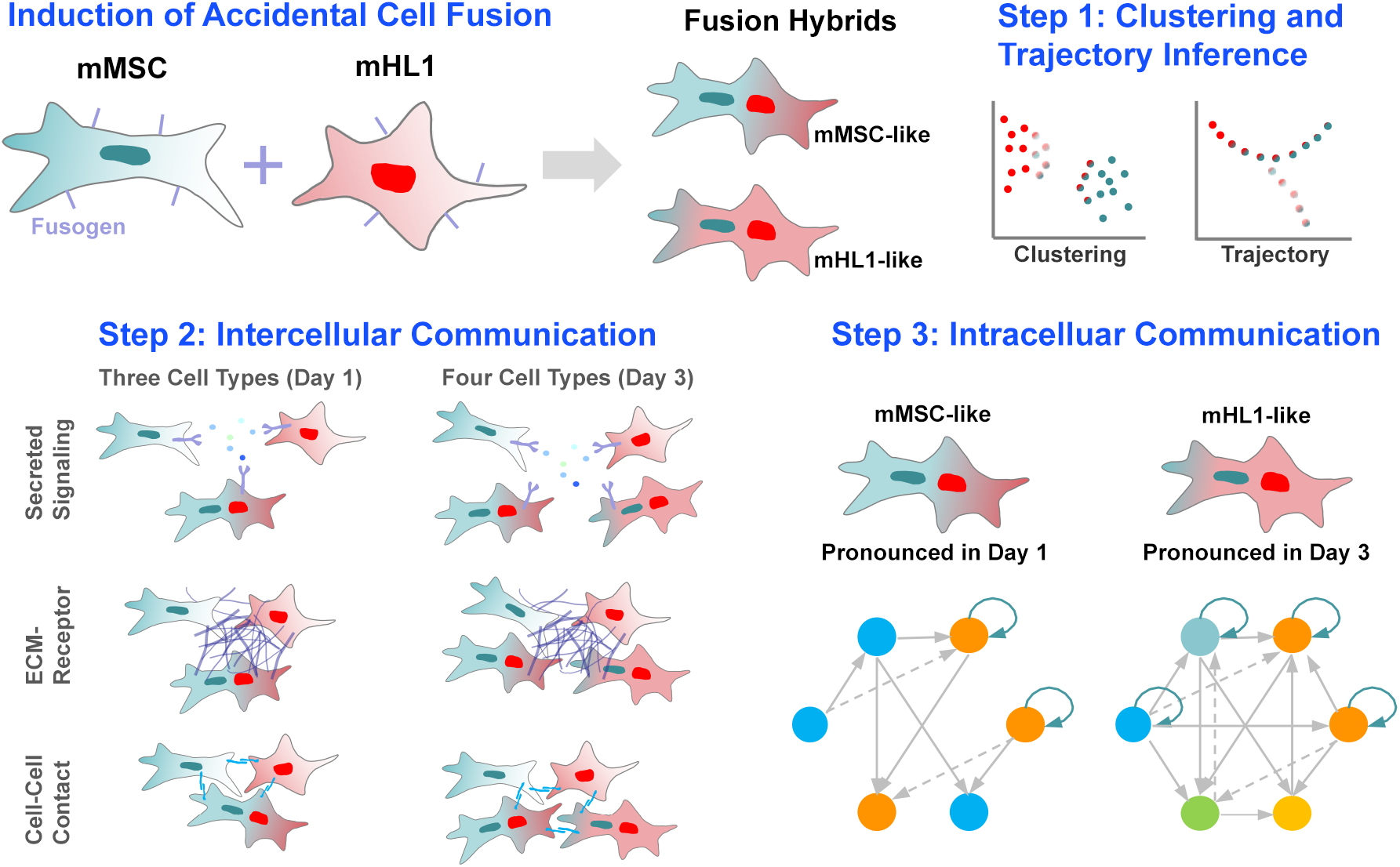
A schematic diagram outlining the experimental design and analytical approach employed in this study. The investigation comprised three key phases: (1) Unsupervised clustering and TI were applied to the co-culture scRNA-seq datasets. (2) Intercellular communication patterns were analyzed to compare interactions between mMSC, mHL1, and their fused hybrid cell populations at both Day 1 and Day 3. (3) Intracellular gene regulatory networks were inferred to characterize the transcriptional reprogramming events associated with cell fusion across the Day 1 and Day 3 time points.

## Methods

### Data acquisition and samples

The original FPKM gene expression matrix obtained from the Gene Expression Omnibus (GEO, accession number GSE69926) was converted to TPM for subsequent analysis. TPM is generally considered more robust and preferred for most analyses, especially when comparing gene expression across different samples^24^. For pre-fusion cells, parental controls consisted of 15 cells of each parental type (mMSC_1–15 and mHL1_1–15) isolated prior to co-culture. After fusion, each cell type was harvested and subjected to scRNA-seq^23^. Fusion cells comprised 28 fusion products identified using BiFC (BiFC_D1_F1–5, 24 h) and dual color (DC) expression of GFP and mCherry (DC_D1_F1–16, 24 h; DC_D3_F1–7, 72 h). Additionally, five cells of each parental cell type (mMSC_D1_1–5 and HL1cm_D1_1–5) exposed to co-culture conditions, but did not fuse were included.

### Marker gene analysis of distinct cell types

To generate AnnData files, we converted TPM values from the available FPKM gene matrix and metadata using Scanpy (ver. 1.10.1) in Python (ver. 3.12.2). We skipped normalization and highly variable gene selection during preprocessing. Instead, we filtered the data by retaining only genes expressed in at least one cell and cells expressing at least 200 genes. Next, we applied a log(x+1) transformation directly to the counts matrix. We then computed the K-nearest neighbors for each cell, using the 17 nearest neighbors in a seven-dimensional principal component space. Finally, we visualized the high-dimensional embedding using Uniform Manifold Approximation and Projection (UMAP) overlaying our known annotations from the metadata.

#### Clustree

To compare our known annotations with unsupervised clustering, we applied Clustree^25^. The raw AnnData was converted into a Seurat object and processed through normalization, feature selection and scaling. Both PCA and UMAP was employed to reduce dimensionality, followed by clustering at multiple resolutions (0.2, 0.4, 0.6, 0.8, 1.0) to identify cell clusters. Furthermore, we visualized Clustree graphs using the clustree function, where resolutions were specified with SNN (shared nearest neighbor) resolution, nodes are clusters, and edges indicate overlap between clusters.

#### FindAllMarkers

Our approach to identifying differentially expressed genes began with the FindAllMarkers function, which compares gene expression in each cluster against all others. We specifically sought upregulated markers, setting a log_2_ fold change threshold at 0.25 and accepting all other default settings. A subsequent filtering step ensured that only highly significant genes were carried forward: markers had to exhibit a log_2_ fold change of 2 or more, a p-value under 0.05, and a delta percent expression greater than 20%. This curated list of markers was then exported to a CSV file for downstream analysis.

#### Venn diagram

We compared the number of differentially expressed gene markers identified by our known annotation and unsupervised clustering. We visualized these results using a Venn diagram, where numbers represent gene counts in each specific cell type and cluster.

### Trajectory Inference

To investigate the transcriptional dynamics of mHL1 and mMSC cells during fusion and understand the developmental trajectory of the resulting hybrids, we utilized the Monocle 3 package^26^. We first established a CellDataSet object, incorporating the UMAP dimensionality reduction previously computed in Seurat to ensure consistency across downstream analyses. Cells were then clustered within Monocle 3 to define cellular states for trajectory inference. We constructed a single trajectory, specifically designed to allow both parental cell types at Day 0 to serve as root cells, enabling the calculation of pseudotime and the ordering of hybrid cells along this developmental continuum. The inferred trajectory and pseudotime progression were visualized on the UMAP embedding. To identify genes significantly associated with this trajectory, we performed differential gene expression (DGE) testing and subsequently grouped these genes into co-expressed modules. Next, we examined the expression patterns of these modules along the trajectory. Representative cell types were then identified using the EnrichR analysis tool^27^ with Tabula Muris atlas^28^.

### Intercellular communications: CellCall and CellChat

We evaluated intercellular communication between cell types using two sets of cells: 31 from Day 1 co-culture and 38, including fusion cells, from Day 3. The latter aimed to provide a broader understanding of how cell-cell communication changes between Day 1 and Day 3. R packages CellCall^29^ and CellChat^30^ in R (v4.3.3) were used for analysis. Both packages leverage known ligand-receptor interactions to construct intercellular communication networks. The initial output of CellCall included a Circos plot visualizing signaling between cell types, indicating ligand and receptor roles. Subsequently, a bubble plot displayed associated pathways for each cell type combination, colored by their normalized enrichment score (NES). CellChat identified major signaling pathways in cell-cell interactions, with the top five to six pathways based on relative contribution being further analyzed. CellChat analysis^31^ typically requires a minimum of 10 cells per cell type^32^. However, due to the limited number of total cells in our analysis, we adjusted this parameter to a minimum of 2 cells per cell type.

### Intracellular communications: Gene regulatory network

We employed the pySCENIC (Single-Cell rEgulatory Network Inference and Clustering) workflow (ver. 0.12.1) on the NIH Anvil Platform to analyze gene regulatory networks (GRNs) and assess regulon activity. Our analysis leveraged cisTarget databases for ranking and scoring, a transcription factor (TF) list and a motif enrichment table. First, we identified co-expressed genes based on enriched DNA motifs within 10 kb or 500 bp of the transcription start site (TSS) using RcisTarget databases. The pySCENIC’s GRNBoost2 algorithm then inferred GRNs to identify co-expression modules. Next, cisTarget motif enrichment analysis was performed on these modules to detect regulons. We used motif rankings and scores to identify TF binding motifs, which were subsequently matched to TFs using the motif enrichment table. We quantified regulon activity using AUCell. This method ranks cells based on their regulon gene expression and calculates the area under the recovery curve (AUC) to assess activity. Following this, we computed regulon specificity scores (RSS) using default settings to gain insights into regulon specificity and activity across different cellular contexts. To identify master regulators within each cell cluster, we transformed the activity scores into binary matrices using the binarize function with default settings. We then extracted master regulators and their gene targets with an importance score greater than two from the adjacencies file generated by the GRNBoost2 function. Finally, the extracted gene list was imported into Cytoscape (v3.10.1) to visualize the predicted intracellular GRN by defining nodes and edges.

## Results

### Annotation and Unsupervised Learning Distinguished Cell Types, and Pseudotime Analysis Revealed Significant Temporal Alterations in Gene Expression

**Figure 2** presents a multi-faceted scRNA-seq analysis, dissecting the distinct transcriptional profiles and trajectories of mHL1 cells, mMSCs, and fusion hybrids. Replicating our prior PCA^23^ (**Figure 2A**), we further employed UMAP (**Figure 2B**) for enhanced dimensionality reduction. UMAP more distinctly resolved the cell populations, primarily segregating them into two major categories. Notably, hybrids with a more mMSC-like transcriptome (mMSC-hybrids) and hybrids with a more mHL1-like transcriptome (mHL1-hybrids) occupied separate territories, partitioned from their non-fused counterparts. This segregation, indicating substantial gene expression alterations upon fusion, was less evident in the PCA. To investigate cell population structure, we performed unsupervised clustering using Clustree, which identified three clusters (**Figure 2C**), contrasting with the four cell types delineated by UMAP. DGE analysis (**Figure 2D** and **2E**) pinpointed key marker genes defining each annotated cell type and unsupervised cluster, underscoring the distinct transcriptional identities revealed by both approaches. This is further visualized in the Venn diagram (**Figure 2F**), quantifying transcript overlap between cell types and clusters. Notably, mHL1 and mHL1-hybrids exhibited considerable transcript overlap with Cluster 0 (290 and 273, respectively), which also displayed a substantial number of uniquely expressed transcripts (455). Conversely, mMSC-hybrids and mMSC cells showed minimal transcript overlap with Cluster 1 and Cluster 2 (two and zero, respectively), prompting the question of whether this disparity reflects changes beyond the inherent biological differences between mMSC fusion and mMSC cells.

**Figure 2.**
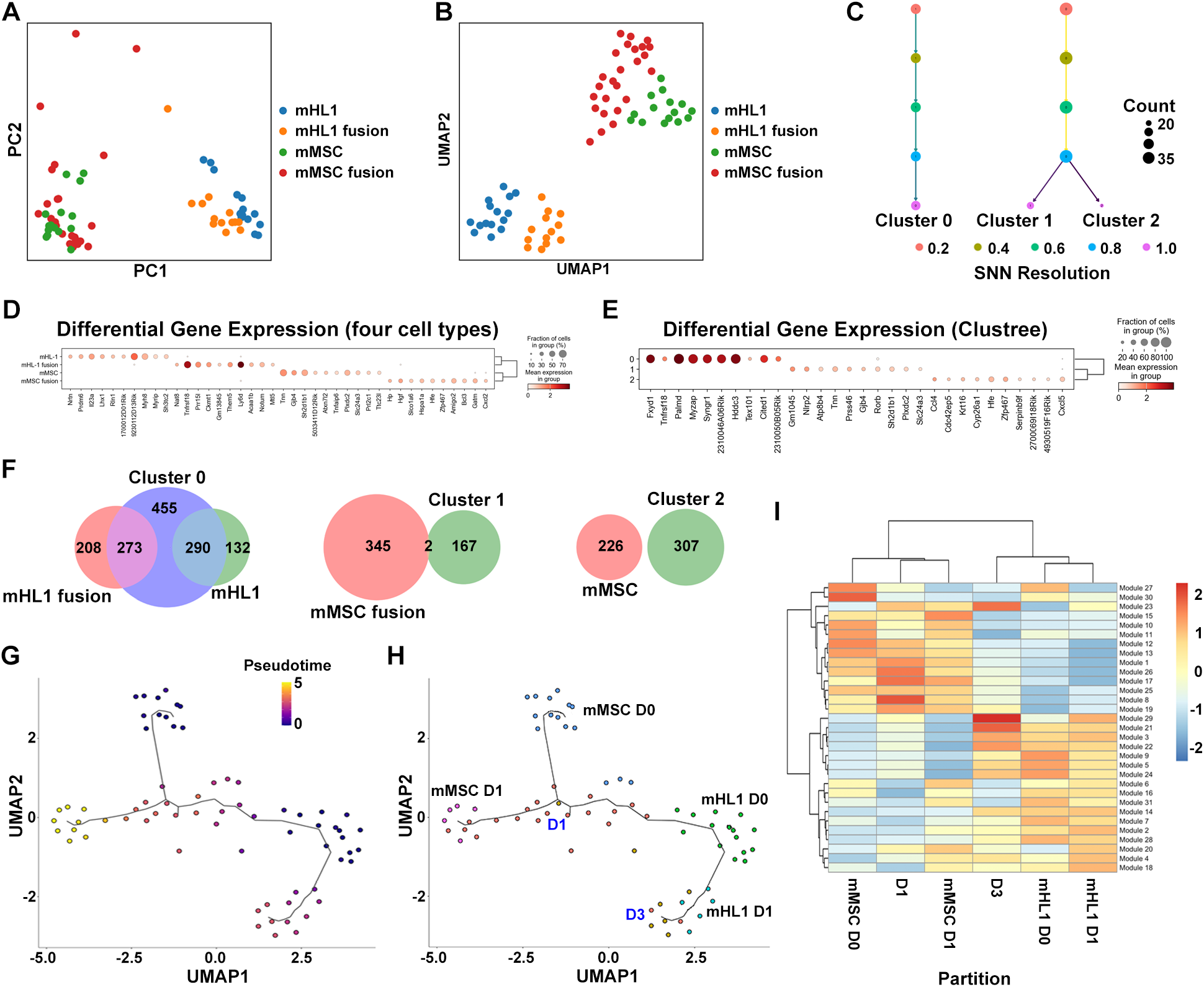
scRNA-seq analysis of mHL1, mMSC, and their fusion products. Cell type separation is visualized in the PCA plot in (A) and the UMAP embedding in (B). Clustree analysis in (C) illustrates cluster stability and relationships across different SNN resolutions. Differentially expressed genes across the four main cell populations and Clustree-identified clusters are shown in the dot plots in (D) and (E), respectively. Venn diagrams in (F) depict the number of unique and overlapping genes in mHL1, mMSC, and their fusion products within Cluster 0, Cluster 1, and Cluster 2. TI analysis is projected onto the UMAP embedding in (G), with cells colored by pseudotime, and in (H), highlighting the inferred differentiation paths of mMSC and mHL1 cells over three days. The heatmap in (I) displays the expression of gene modules across different cell types and time points (D1 and D3) after partitioning.

To explore the temporal dynamics of gene expression, we performed pseudotime analysis to infer transitional transcriptional states, particularly within the fusion products. The pseudotime analysis (**Figure 2G**) effectively resolved the different cell types along a trajectory, where a color gradient from darker (earlier pseudotime) to lighter hues (later pseudotime) ordered cells based on their inferred progression through transcriptional states. Overlaying cell type annotations onto this inferred trajectory (**Figure 2H**) revealed that mMSC fusion and mHL1 fusion cells represented terminal partitions originating from the designated root cells of mMSC and mHL1, respectively. Notably, the inferred pseudotime difference between mMSC D0 and mMSC D1, as indicated by a substantial shift in color from darker to lighter, appeared larger than the corresponding transitions observed in the mHL1 lineage (e.g., from its inferred origin towards mHL1 D1 or D3). This suggests a more extensive degree of inferred transcriptional remodeling occurred within the mMSC lineage (hybrids and unfused mMSCs) during this pseudotime interval compared to the mHL1 lineage. Finally, the heatmap (**Figure 2I**) visualized global gene expression patterns by displaying modules of co-regulated genes and their differential expression across cell types and pseudotime. This analysis corroborated the transcriptional divergence of the fusion-derived populations, providing a complementary perspective to the trajectory analysis in **Figures 2G** and **2H**. The observed larger pseudotime difference between mMSC D0 and D1 in the UMAP underscores a more pronounced transcriptional shift during this transition compared to the visualized changes within the mHL1 lineage along the inferred pseudotime.

While the UMAP visualization and overlaid trajectory captured the overall fusion landscape and individual cell ordering, the heatmap (**Figure 2I**), by averaging gene module expression within labeled groups, offered a consistent view of the relative pseudotime? evolution? of these populations and further highlighted the distinct transcriptional states of the fusion products. The top five cell types identified by Tabula Muris based on the transcriptional state for each module are presented in **Table S1**. Analysis of Monocle 3 modules in mMSC and mHL1 cells reveals dynamic cell type changes across partitions. Early mMSC shows Fibroblasts (Heart) and Mesenchymal cells (bladder), transitioning to other cell types and later featuring Stromal Cells (Trachea). Early mHL1 is dominated by Stem Cells of Epidermis (Skin) and Cardiac Muscle Cells, later including Keratinocytes (Tongue). The recurrence of Cardiac Muscle Cells in mHL1 and shared later cell types like Stromal Cells (Trachea) suggest potential convergence. Overall, these dynamic changes in cell type prevalence underscore the complex cellular transitions and heterogeneity enabled by cell fusion, with the tissue of origin providing valuable context for their potential roles.

### Distinguished Patterns of Intercellular Signaling Were Identified

To gain insights into the intercellular communication landscape of the Day 1 cells (n=31), we utilized Circos plots (**Figures 3A** and **3B**) to visualize the relationships between cell groups defined by annotation and Clustree analysis. Unsupervised clustering of these 31 cells independently identified three distinct cell types (**Figure S1A**). Notably, the number and interconnectedness of clusters revealed by Clustree diverged from the pre-defined annotated cell types (Fusion, mHL1, mMSC). This discrepancy suggests that unsupervised learning may be capturing additional layers of cellular heterogeneity or grouping cells based on unique transcriptional signatures not fully represented by the initial annotation.

**Figure 3.**
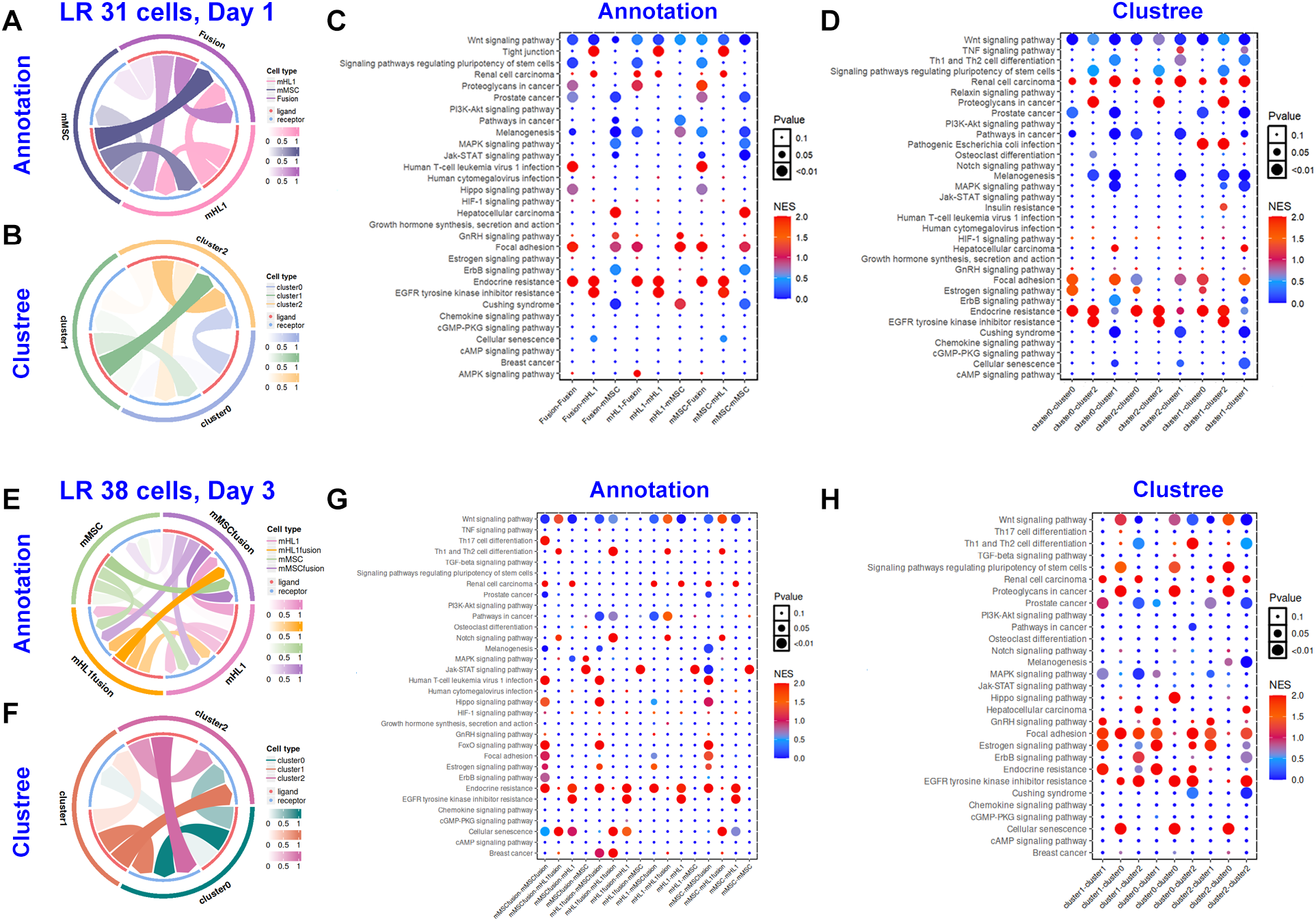
LR interactions are visualized as Circos plots, organized by cell type annotation (A, E) and Clustree clustering (B, F). Dot plots show enriched signaling pathways based on cell type annotation (C, G) and Clustree clustering (D, H). Dot size indicates pathway significance, and color represents the NES. Data for Day 1 (n=31 cells) is shown in panels A-D, and data for Day 3 (n=38 cells) is shown in panels E-H.

Dot plots in **Figures 3C** and **3D** visualize the enriched pathways identified. Comparative pathway enrichment analysis between annotation-defined cell types and Clustree-identified clusters revealed similarities and differences in the transcriptional landscape of Day 1 cells. In **Figure 3C**, pathways with a p-value ≤ 0.05 and at least 5 cell type pairs (representing >50% of permutations) showed higher NES for Focal adhesion (6 pairs) and Endocrine resistance (6 pairs), while Wnt signaling (9 pairs) and Melanogenesis (6 pairs) exhibited lower NES values. Downregulation of Wnt signaling suggests a reduced role in development and proliferation within these communications. Similarly, decreased melanogenesis hints at fewer related functions like antioxidant defense, a role that is relevant to MSCs through potential indirect mechanisms such as interaction with melanin from other cells or shared pathway components^33^. Conversely, upregulated Focal adhesion indicates increased emphasis on physical cell interactions, while upregulated Endocrine resistance suggests reduced hormonal sensitivity. Within Clustree-identified cell types (**Figure 3D**), Renal cell carcinoma (9 pairs) displayed higher NES, whereas Wnt signaling (9 pairs), Pathways in cancer (5 pairs), and Melanogenesis (5 pairs) showed lower NES. Notably, Endocrine resistance (9 pairs) and Focal adhesion (6 pairs) presented mixed NES values across cell type pairs. The consistent upregulation of Renal cell carcinoma pathways across all cell-cell communication pairs suggests the utilization of signaling mechanisms common in kidney cancer, while the downregulation of Wnt signaling across all pairs points to a reduced role in developmental and proliferative communication. The mixed NES values for Endocrine resistance and Focal adhesion indicate context-dependent influences of these pathways on cell-cell interactions. Overall, this pattern reveals a communication landscape characterized by the prominence of Renal cell carcinoma-related signaling and a diminished role for Wnt, with the importance of endocrine resistance and physical interactions varying between specific cell type pairs.

Extending the cell culture period by two days (n=38) enabled the identification of four distinct cell types through annotation (**Figure 3E**). In contrast, Clustree analysis consistently identified three cell types (**Figure 3F** and **S1B**). Day 3 cell-cell communication revealed distinct pathway patterns. No single pathway encompassed all 16 annotations (**Figure 3G**) or nine Clustree groups (**Figure 3H**). In **Figure 3G**, pathways exhibiting a p-value ≤ 0.05 and present in over 50% of cell type pair permutations showed elevated NES for Endocrine resistance (8 pairs) and mixed NES values for Wnt signaling pathways (11 pairs). The distinct pathway patterns observed in Day 3 cell-cell communication suggest a complex and multifaceted communication network. Specifically, the consistent upregulation of Endocrine resistance pathways across a majority of cell type interactions in **Figure 3G** implies a significant and widespread influence on how these cells might respond to hormonal signals during their communication. The mixed NES values for Wnt signaling in the same analysis indicate a more context-dependent role for this crucial developmental pathway, suggesting it might be activated in some communication axes but not others, highlighting heterogeneity in the signaling mechanisms employed by Day 3 cells. This initial snapshot reveals a communication landscape where resistance to endocrine cues and a variable engagement of Wnt signaling are prominent features. **Figure 3H** highlighted higher NES values for Focal adhesion (7 pairs) and EGFR tyrosine kinase inhibitor resistance (5 pairs), alongside mixed NES values in Wnt signaling (7 pairs), Estrogen signaling (6 pairs), and Prostate cancer (5 pairs) pathways. **Figure 3H** reveals a different facet of Day 3 cell-cell communication, characterized by a consistent upregulation of Focal adhesion pathways, suggesting a significant role for physical cell interactions and mechanotransduction^34^ in these communication axes. Similarly, the elevated NES for EGFR tyrosine kinase inhibitor resistance implies that mechanisms associated with resistance to this cancer therapy are also prominent in the signaling between these cell types. The mixed NES values observed for Wnt signaling, Estrogen signaling, and Prostate cancer pathways indicate that the influence of these pathways on cell-cell communication is context-dependent, being more active in certain cell pairings than others, thus highlighting the heterogeneity and specificity of signaling within the Day 3 cell population.

The full spectrum of intercellular communication among 31 or 38 co-cultured cells, classified by either annotated or Clustree-identified cell types, is illustrated in **Figure S2**. A more in-depth analysis of specific signaling pathways across three different modes of intercellular communication follows in the next section.

### Distinct Modes of Intercellular Communication, Active Across All Cell Types, Are a Key Feature of the Observed Cellular Interactions

For each of the three intercellular communication modes, we identified the top five most significant pathways, with the exception that all LR pairs are included within four pathways in **Figure 4B**. The complete breakdown of the relative contributions from all signaling pathways is presented in the supplementary material: **Figures S3** and **S4** for the 31 cells (Day 1), and **Figures S5** and **S6** for the 38 cells (Day 3).

**Figure 4.**
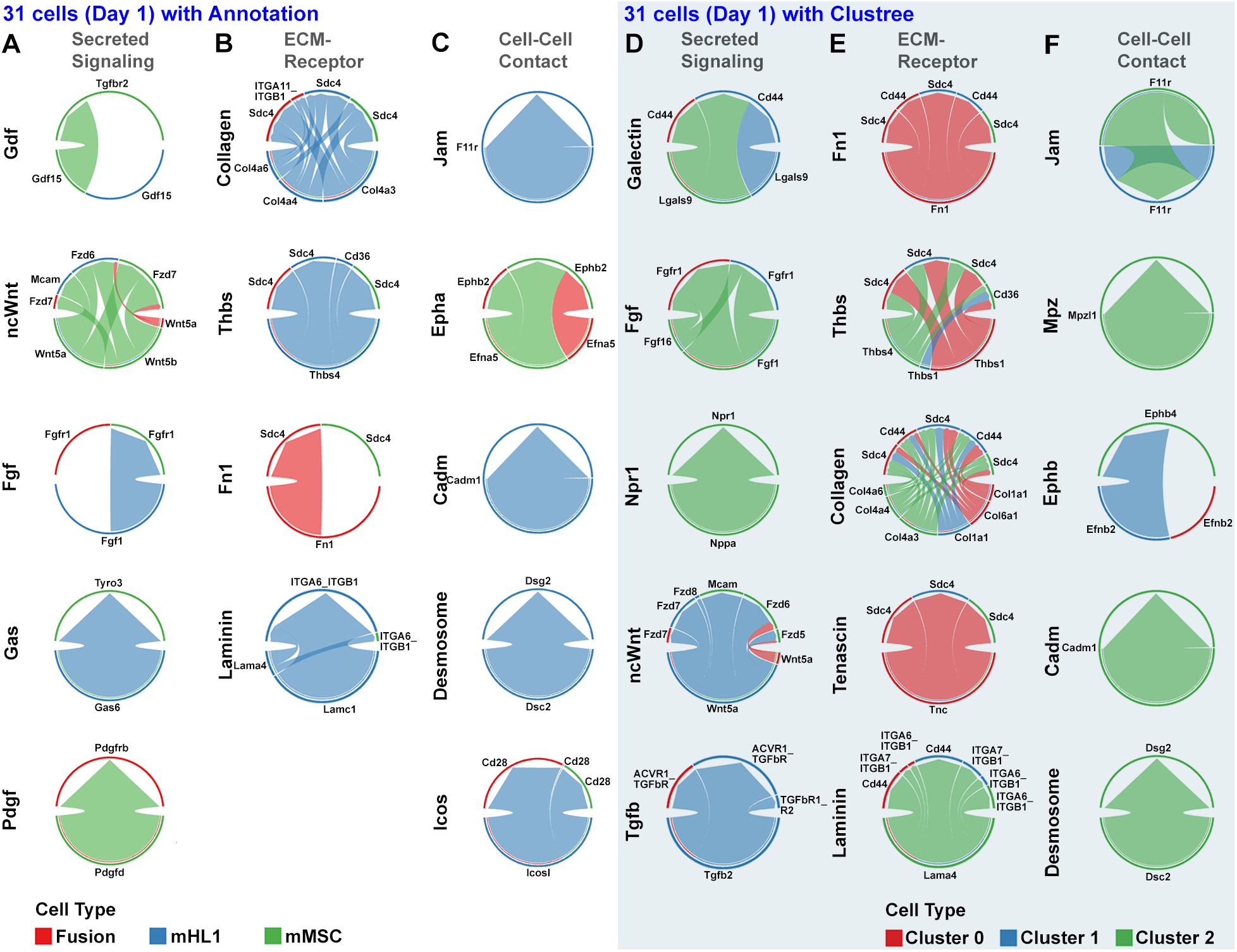
LR pair contributions to cell-cell communication in 31 co-cultured cells at Day 1, analyzed by communication type and cell identity. In (A-C), contributions based on cell type annotation (Fusion, mHL1, mMSC) are shown for Secreted Signaling, ECM-Receptor interactions, and Cell-Cell Contact, respectively. In (D-F), corresponding contributions based on unsupervised clustering (Cluster 0, Cluster 1, Cluster 2) are presented. Each chord diagram illustrates the relative contribution of specific LR pairs within each communication type, with segment size proportional to the contribution score and colors indicating the sending and receiving cell populations. Note: Gas (G protein subunit alpha s).

Analysis of cell-cell communication using both cell type annotation and unsupervised clustering with Clustree yielded distinct insights into the cell-cell communication landscape of the 31 cells at Day 1. Based on the annotated cell types (**Figure 4A**), mMSC cells expressed LR pairs associated with Gdf (Growth Differentiation Factor) pathways, a subset of the TGFβ (Transforming Growth Factor-β) superfamily, while Fusion cells exhibited more localized signaling within the Fgf (Fibroblast Growth Factor) and Pdgf (Platelet-Derived Growth Factor) pathways. Notably, while Gdf signaling was prominent in the annotated cell types, the Clustree analysis indicated a key role for TGFβ pathways in the identified clusters. However, Clustree-based clustering (**Figure 4D**) revealed a more nuanced perspective, with Cluster 0 demonstrating an enrichment in Galectin signaling pathways, a characteristic not evident from the annotated cell types. Furthermore, ncWnt (non-canonical Wnt) signaling displayed heterogeneity across both the annotated cell types and the Clustree-defined clusters, suggesting potential sub-population specific signaling activities. The main differences in terms of related pathways between the annotation-based set (Gdf, ncWnt, Fgf, Gas, Pdgf) and the Clustree-identified cell types set (Galectin, Fgf, Npr1, ncWnt, TGFb) lie in their mechanisms of action and primary roles. The annotation-based set primarily features receptor tyrosine kinase (RTK) and serine/threonine kinase signaling for growth factors and G protein-coupled receptor (GPCR) signaling via Gas (G protein subunit alpha s)^35^. In contrast, the Clustree-identified cell types set, while sharing Fgf (RTK) and ncWnt, introduces glycan-mediated interactions through Galectins and guanylate cyclase-linked receptor signaling via Npr1^36,37^, alongside the serine/threonine kinase signaling of the TGFβ superfamily. This shift highlights a potential emphasis in the early co-culture dynamics, as revealed by the Clustree analysis, on cell-cell and cell-matrix interactions and fluid homeostasis/cGMP signaling compared to the more classical growth factor and broad GPCR signaling expectations based on cell type annotation.

Examining ECM-receptor interactions revealed a predominant expression of Collagen and its receptors in the mHL1 annotated cell type (**Figure 4B**), suggesting a key role for these interactions in this specific population. However, Clustree-based clustering (**Figure 4E**) uncovered a more diverse interaction profile across all Clustree-identified clusters, characterized by strong connections between specific Collagen subtypes and Sdc4(Syndecan-4)/CD44 receptors, potentially reflecting distinct engagement with multiple LR pairs within these subpopulations. Similarly, while Thbs (Thrombospondin) signaling pathways appeared dominant in mHL1 cells based on annotation, Clustree analysis indicated involvement of all three clusters in these pathways. For Fn1 (Fibronectin1) and Laminin signaling, a primary association with a single cell type was observed in both annotation and Clustree. Nevertheless, the overall trend of increased diversity in ECM-receptor interactions revealed by Clustree analysis (**Figure 4E**) highlights its capacity to identify finer-grained distinctions. The prominence of Tenascin and its corresponding receptors further underscores the ability of Clustree to identify subpopulations with unique ECM interaction signatures. The annotation-based ECM component list (Collagen, Thbs, Fn1, Laminin) represents expected key structural and matricellular components based on the anticipated ECM of the involved cell types. The Clustree-identified list (Fn1, Thbs, Collagen, Tenascin, Laminin) largely overlaps but uniquely highlights Tenascin, suggesting that specific cell populations within the co-culture are expressing this matricellular protein associated with dynamic tissue remodeling and complex effects on cell behavior^38^. While both lists point to the importance of core ECM components like Collagen, Fibronectin, and Laminin, the unsupervised learning approach provides a more specific, data-driven insight into the ECM landscape by identifying Tenascin as a potentially significant and differentially expressed factor within the co-culture at this time point.

Finally, the analysis of cell-cell contact molecule expression revealed notable differences between the cell type annotation and Clustree-based clustering. For instance, Jam (Junctional Adhesion Molecule) signaling, which appeared primarily driven by mHL1 through a single LR pair in the annotated cell types (**Figure 4C**), was associated with two clusters in the Clustree analysis (**Figure 4F**). Similarly, while Epha (Eph receptor A) signaling was prominent in the annotated cell types, Clustree analysis highlighted the significance of Ephb (Eph receptor B) signaling pathways. Conversely, the expression patterns of both Cadm (Cell Adhesion Molecule) family members and desmosome-related components showed considerable consistency across both analytical methods. Interestingly, Icos (Inducible T-cell Costimulator) pathways were mainly linked to mHL1 in the annotated cell types, whereas Mpz (Myelin Protein Zero) pathways were a notable feature of Cluster 2 in the Clustree analysis. The annotation-based Cell-Cell contact pathway list (Jam, Epha, Cadm, Desmosome, Icos) reflects expected adhesion and interaction mechanisms based on the anticipated cell types, including T cell costimulation via Icos. The Clustree-based list (Jam, Mpz, Ephb, Cadm, Desmosome) reveals a slightly different emphasis. While sharing core adhesion pathways like Jam, Cadm, and Desmosome, it highlights Ephb instead of Epha for Eph/ephrin signaling, suggesting a preference for B-type ephrin interactions in the identified cell clusters. Notably, the Clustree analysis also uniquely identifies Mpz, a myelin-specific adhesion molecule^39^, potentially indicating the presence of unexpected cell types or a novel function for Mpz in this co-culture context, while Icos, associated with T cell interactions^40^, is not prominent in the Clustree-defined clusters at this stage.

Following two additional days of co-culture, the intercellular communication networks of 38 cells were investigated and contrasted using two distinct cell classification approaches: annotation-based identification of four cell types (mHL1, mHL1 fusion, mMSC, and mMSC fusion) and unsupervised clustering via Clustree, which delineated three distinct clusters (Cluster 0, Cluster 1, and Cluster 2). Analysis of Secreted Signaling pathways revealed shared key pathways mediating communication within and between both the annotated cell types (**Figure 5A**) and the Clustree-defined clusters (**Figure 5D**), including Galectin, Spp1 (Secreted Phosphoprotein 1), Angptl (Angiopoietin-like), and Fgf. Notably, Mif (Macrophage Migration Inhibitory Factor) signaling was prominent in the annotated cell type analysis, whereas Gas signaling appeared more influential in the Clustree-identified cell types. Furthermore, regarding Spp1 signaling, mMSC and mMSC fusion cells exhibited a greater diversity of integrin receptors compared to the cell types defined by Clustree. The annotation-based secreted signaling pathway list (Galectin, Spp1, Angptl, Mif, Fgf) reflects expected secreted factors involved in cell interactions, angiogenesis, and inflammation based on the anticipated cell types. The Clustree-based list (Galectin, Spp1, Fgf, Angptl, Gas), shows a strong overlap in key factors like Galectin, Spp1, Angptl, and Fgf, indicating their continued importance. However, the annotation-based list highlights Mif, suggesting a potential role for macrophage migration inhibitory factor in the expected signaling^41^, while the Clustree analysis uniquely identifies Gas, indicating a prominent role for GPCR signaling pathways linked to Gas^42^ in the specific cell populations defined by unsupervised learning at this later time point.

**Figure 5.**
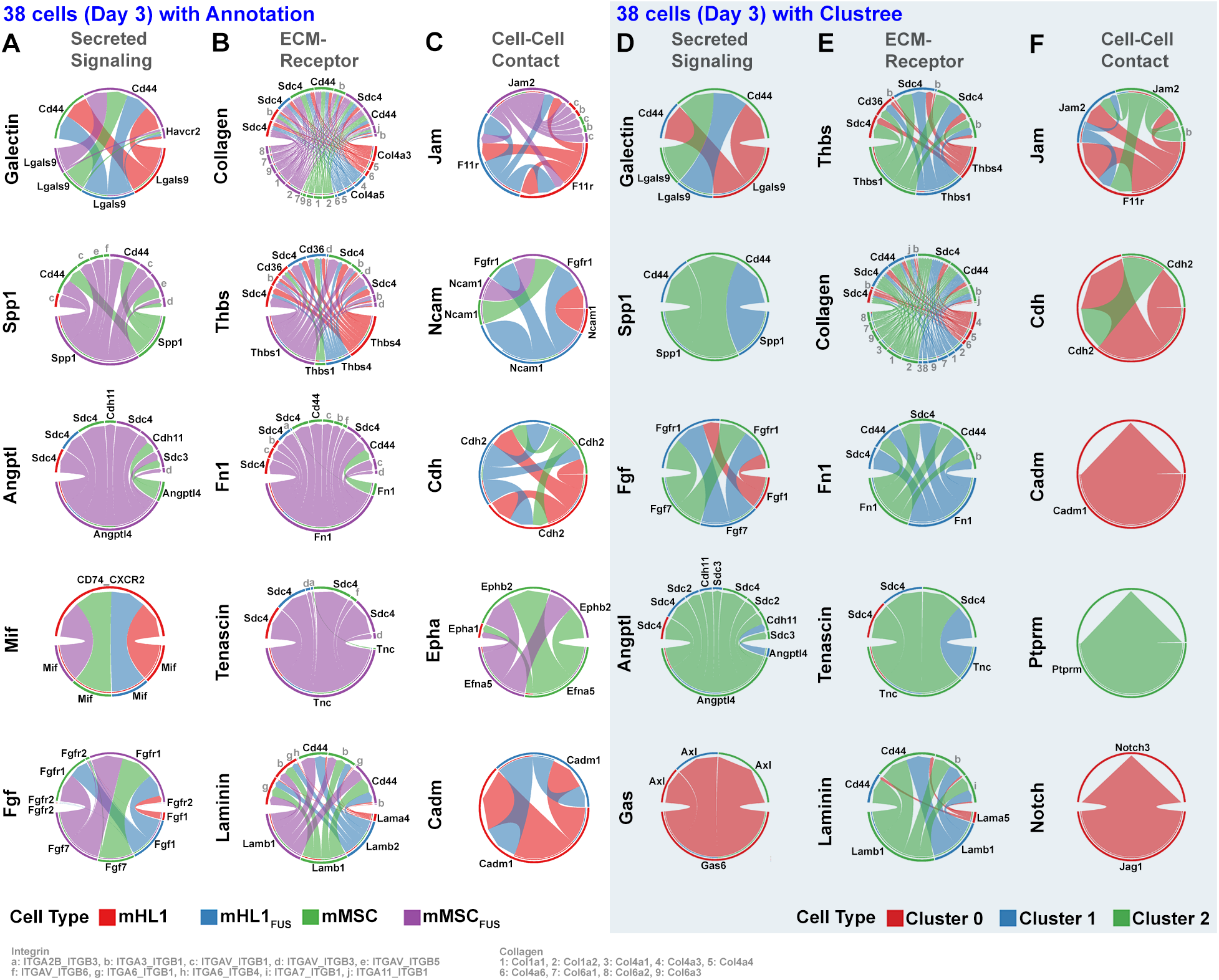
LR pair contributions to cell-cell communication in 38 co-cultured cells at Day 3, analyzed by communication type and cell identity. In (A-C), contributions based on cell type annotation (mHL1, mMSC, mHL1 fusion, mMSC fusion) are shown for Secreted Signaling, ECM-Receptor interactions, and Cell-Cell Contact, respectively. In (D-F), corresponding contributions based on unsupervised clustering (Cluster 0, Cluster 1, Cluster 2) are presented. Each chord diagram illustrates the relative contribution of specific LR pairs within each communication type, with segment size proportional to the contribution score and colors indicating the sending and receiving cell populations. To enhance clarity, specific integrin subunits (a-j) and collagen isoforms (1-9) are listed separately.

Similarly, the analysis of ECM-Receptor interactions (**Figures 5B** and **5E**) revealed the involvement of shared ECM pathways across both cell classification methods, albeit with varying extents of LR pair representation within the different cell types/clusters. To manage complexity, integrin subunits were encoded from ‘a’ to ‘j’, and collagens from ‘1’ to ‘9’. In general, the annotation-based approach with four cell types identified a more complex repertoire of LR pairs compared to the Clustree-based analysis with three clusters. Integrin involvement was a consistent feature across all four annotated cell types, a characteristic that was notably absent in the Tenascin interactions observed within the Clustree-identified cell types (**Figure 5E**). The annotation-based and Clustree-identified lists of ECM components (Collagen, Thbs, Fn1, Tenascin, Laminin) in the Day 3 co-culture analysis are remarkably identical, differing only in their order. This strong agreement indicates that the major ECM components expected based on the annotated cell types are indeed being highlighted by the unsupervised learning analysis of the gene expression data. These five components likely represent the core ECM environment in the co-culture at this stage, encompassing structural support and dynamic regulatory factors, and suggesting that the anticipated ECM landscape is well-reflected in the molecular data.

Finally, the analysis of Cell-Cell Contact interactions (**Figures 5C** and **5F**) reveals the key adhesion molecules mediating direct cell-cell communication through homotypic or heterotypic interactions. Comparison of these panels highlights distinct profiles between the annotation-based cell types and the Clustree-derived clusters, with the latter exhibiting a less complex repertoire of LR pairs (**Figure 5F**). Despite these differences, Jam, Cdh (Cadherin), and Cadm interactions were common to both classification methods, albeit with varying degrees of complexity. Ncam (Neural Cell Adhesion Molecule) and Epha interactions were observed in the annotated cell types, whereas Ptprm (Protein Tyrosine Phosphatase Receptor Type M) and Notch interactions appeared specifically within the Clustree-identified cell types, each characterized by a single LR pair on one cell cluster. The annotation-based list of secreted signaling pathways (Jam, Ncam, Cdh, Epha, Cadm) reflects expected cell adhesion-related molecules with potential soluble signaling roles based on the anticipated cell types. The Clustree-identified list (Jam, Cdh, Cadm, Ptprm, Notch), while sharing Jam, Cdh, and Cadm, diverges by highlighting Ptprm, a protein tyrosine phosphatase involved in adhesion and guidance^43^, instead of Ncam, and Notch, a key developmental signaling pathway involving receptor cleavage and transcriptional regulation^44^, instead of Epha, a RTK primarily known for cell-cell contact-dependent boundary formation^45^. This suggests that the unsupervised learning approach identifies a signaling landscape with a potentially greater emphasis on phosphatase activity (Ptprm) and direct transcriptional regulation via cleaved receptors (Notch) compared to the expected roles of Ncam and Epha based on cell type annotation.

### Gene Regulatory Network Analysis Uncovers Intracellular Master Regulators of mMSC and mHL1 Cell Fusion

Here, we delve into intracellular communications and associated TFs. While SCENIC analyzes data per cell, we performed UMAP of 31 (**Figure 6A**) and 38 (**Figure 6D**) cells for visualization. GRNs were constructed by SCENIC, and the master regulators were identified for both 31 (**Figure 6B**) and 38 (**Figure 6E**) cells. The Day 1 GRN (31 cells) reveals distinct subnetworks centered around specific TFs such as *Runx2*, potentially regulating genes involved in early developmental processes; *Prdm4*, possibly influencing chromatin remodeling; *Tbx4*, a known regulator of limb bud development; *Sox6*, implicated in chondrogenesis; *Foxm1*, associated with cell proliferation; *Foxr1*, involved in mesoderm development; *Mef2d*, a key regulator of muscle differentiation; *Tcf15*, a basic helix-loop-helix TF; *Rara*, a retinoic acid receptor; *Rbak*, a transcriptional repressor; *Mybl1*, a proto-oncogene; and *Rarg*, another retinoic acid receptor. In contrast, the Day 3 GRN (38 cells) exhibits a more interconnected structure, with subnetworks around TFs like *Arntl* (BMAL1, Basic helix-loop-helix ARNT-like 1), a core circadian rhythm regulator; *Zfp354c* (also known as CGGBP1, CGG triplet repeat-binding protein 1), involved in DNA binding; *Klf10*, a Kruppel-like factor; *Zfp438*, a zinc finger protein; *Zfp82*, another zinc finger protein; *Hoxa9*, a homeobox gene involved in development; *Rarg* (again observed on Day 1, suggesting its sustained importance); *Fli1*, an ETS family TF; *Hmga2*, a high mobility group protein associated with chromatin structure; *Prrx1*, a paired-related homeobox protein; and the complex *Nr1h3* (Nuclear receptor subfamily 1 group H member 3)/*Pou2f3* (POU domain, class 2, TF 3), indicating potential combinatorial regulation. The increased connectivity on Day 3 suggests a more integrated and potentially coordinated regulatory landscape as the co-culture evolves.

**Figure 6.**
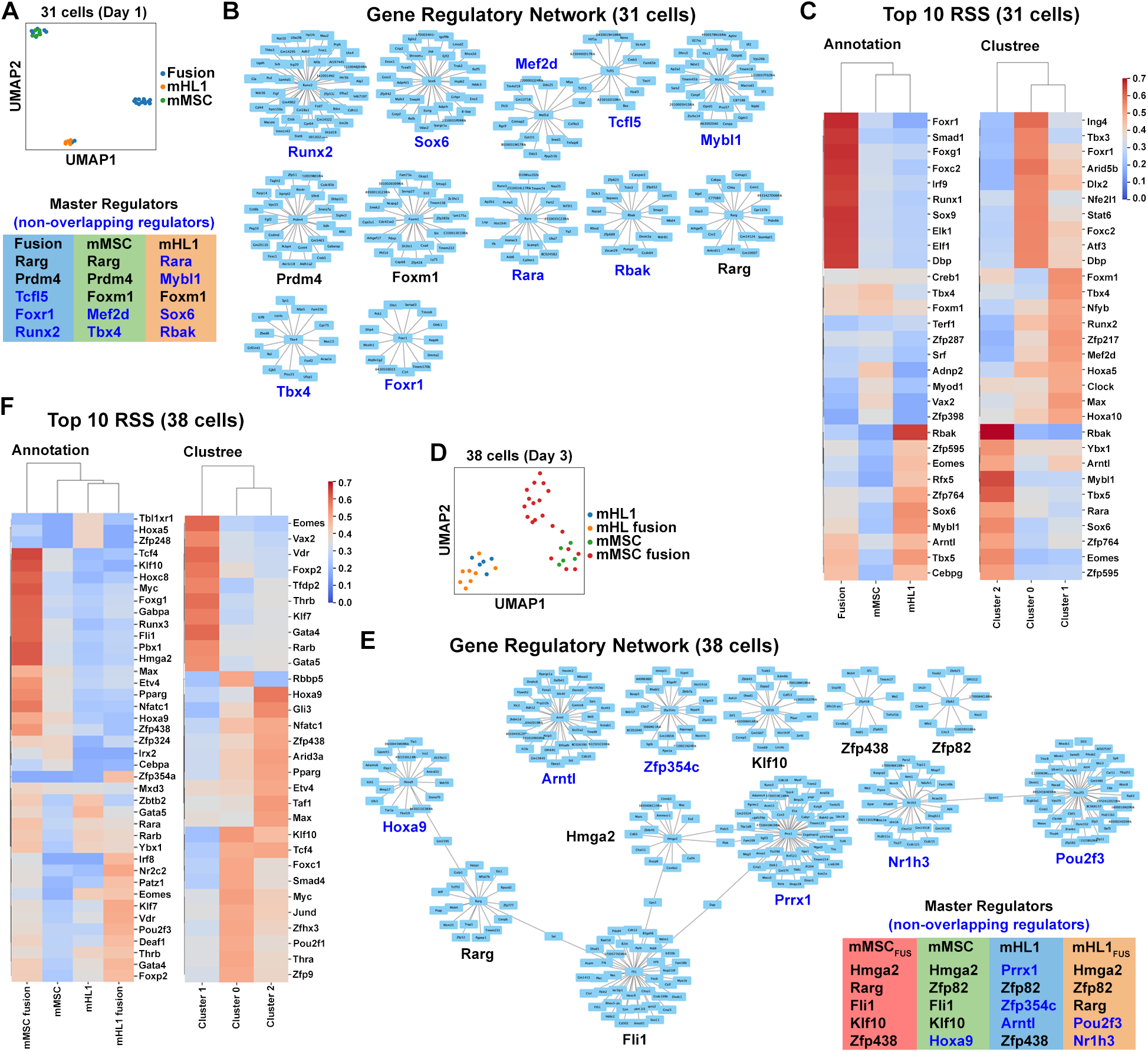
GRN analysis of co-cultured cells at Day 1 (A-C) and Day 3 (D-F). (A) UMAP visualization of Fusion, mMSC, and mHL1 cells at Day 1, with a list of non-overlapping master regulator TFs identified for each. (B) GRNs for selected master regulator TFs at Day 1. (C) Heatmap showing the top 10 RSS for master regulator TFs in Fusion, mMSC, and mHL1 cells (based on annotation) and in Clustree-defined clusters at Day 1. (D) UMAP visualization of mHL1 fusion, mMSC fusion, mHL1, and mMSC cells at Day 3, with a list of non-overlapping master regulator TFs. (E) GRNs for selected master regulator TFs at Day 3. (F) Heatmap showing the top 10 RSS for master regulator TFs in the four cell populations (based on annotation) and in Clustree-defined clusters at Day 3.

The identified master regulators for Day 1 highlight cell-type specific regulatory influences: Fusion cells are characterized by TFs like *Rarg* (involved in retinoid signaling), *Prdm4, Tcf15* (influencing cell fate decisions), *Foxr1*, and *Runx2* (suggesting early developmental programs). mMSCs show enrichment for *Rarg, Prdm4, Foxm1* (indicating proliferative capacity), *Mef2d* (reflecting early differentiation tendencies), and *Tbx4*. mHL1 cells exhibit master regulation by *Rara* (another retinoid receptor), *Mybl1, Foxm1, Sox6* (suggesting chondrogenic potential within this population), and *Rbak*. The presence of shared regulators like *Rarg* and *Foxm1* across different cell groups implies potential common regulatory mechanisms. By Day 3, the master regulator landscape shifts: mMSC fusion cells are defined by *Hmga2* (suggesting chromatin remodeling), *Rarg, Fli1* (involved in cell adhesion and hematopoiesis-related pathways), *Klf10*, and *Zfp438*. mMSCs are characterized by *Hmga2, Zfp82, Fli1, Klf10*, and *Hoxa9* (implying developmental regulation). mHL1 cells show master regulation by *Prrx1* (involved in mesenchymal differentiation), *Zfp82, Zfp354c, Arntl* (suggesting the onset of circadian regulation), and *Zfp438*. Finally, mHL1 fusion cells are marked by *Hmga2, Zfp82, Rarg, Pou2f3* (a POU domain TF), and *Nr1h3* (LXR alpha, a nuclear receptor). The emergence of new key regulators on Day 3, alongside the diminished prominence of some Day 1 regulators, underscores a dynamic reprogramming of transcriptional control within the co-culture.

In the context of single-cell studies, a regulon is defined by a TF and its direct target genes. Ranking regulon activity (RSS, regulon specificity score) across cells provides crucial insights into the dynamic influence of TFs during cell type specification and co-culture experiments. The heatmap in **Figure 6C** displays the top 10 RSS for Day 1, revealing distinct activity patterns. For example, the Runx2 regulon shows high activity in a specific cluster potentially representing early differentiating cells, while the Sox6 regulon is enriched in another cluster possibly indicative of cells with chondrogenic potential. Similarly, the Mef2d regulon is highly active in a subset of cells within the Fusion group, hinting at early muscle-related gene expression programs. Conversely, **Figure 6F** presents the top 10 RSS for Day 3, likely showcasing a different set of influential regulons. For instance, we observe a highly active Hmga2 regulon across several cell types, reflecting its broad role in chromatin remodeling during this later stage. The Arntl regulon becomes more prominent in specific clusters, suggesting the activation of circadian clock-related genes. Furthermore, the activity of regulons like Fli1 or Prrx1 are enriched in particular cell populations, reflecting their roles in processes relevant to the evolving co-culture dynamics. The changes in the top 10 RSS and their enrichment patterns between Day 1 and Day 3 highlight the temporal shifts in key transcriptional drivers and the associated biological processes within the co-culture.

To ensure a comprehensive analysis, we examined non-cocultured parental cells both individually (n=15 per group, **Figure S7**) and collectively (n=30, **Figure S8**). The presence of Hmga2 (**Figure 6E**) in mMSCs and their fusion progeny (mMSC fusion and mHL1 fusion) suggests the inheritance of certain TFs that could function as master regulators in the fused cells. However, our data indicate that the majority of master regulators in the progeny (visualized on UMAPs, **Figure S9**) arose *de novo* rather than being directly inherited from the parental cells.

## Discussion

This study embarked on a detailed characterization of the diverse cell types that emerge from the accidental fusion of mHL1 and mMSC. Our central hypothesis posited that such fusion events would not result in a simple averaging of parental characteristics but would instead rapidly induce the formation of novel cellular phenotypes. We specifically anticipated a biased redistribution of gene expression and cellular mass within the fused cells, suggesting a dominant contribution from one parental cell type over the other. Furthermore, we proposed that the kinetics and extent of these phenotypic changes would be dynamically modulated by a complex interplay of environmental cues and the intrinsic biological attributes of the originating mHL1 and mMSC cells, as well as their hybrid progeny, ultimately leading to temporal variations in gene expression profiles. Given the logistical constraints that precluded a direct, high-resolution time-course experiment, we strategically employed pseudotime analysis, a computational approach that infers the temporal order of cellular states from scRNA-seq data, to dissect these dynamic transcriptional landscapes. Our analysis, visually represented in **Figure 2G-2I**, revealed a more pronounced and protracted alteration in gene expression programs within mMSC fusion products (positioned later along the inferred pseudotime trajectory) compared to their mHL1 counterparts. This observation was further substantiated by the shorter pseudotime durations observed for mHL1 fusion products at both Day 1 and Day 3 post-fusion, suggesting either a faster stabilization of their transcriptional state or a more limited overall extent of change following the fusion event. Consistent with this trend, we identified a greater degree of overlap in gene expression signatures between cells associated with mHL1 fusion and Cluster 0, implying a closer transcriptional relationship. Conversely, cell types associated with mMSC fusion exhibited minimal overlap in gene expression within Clusters 1 and 2 (**Figure 2F**), hinting at the emergence of more distinct transcriptional programs in these hybrid cells.

Building upon the observed bias in gene expression and cell mass, we further hypothesized that this asymmetry would be reflected in the intricate patterns of both intercellular and intracellular communication within the fused cell populations. Specifically, we reasoned that the dominant parental contribution to gene expression would likely shape the signaling pathways and LR interactions that govern how these hybrid cells interact with their microenvironment and regulate their internal processes. To gain a clearer understanding of these dynamic communication changes, we leveraged unsupervised clustering algorithms to group cells based on their transcriptional similarities, aiming to identify distinct cell types emerging post-fusion. Recognizing the inherent challenges in relying solely on pre-defined lineage-specific markers for accurate cell type identification in these novel hybrid entities, we implemented a comparative approach. We evaluated cell type assignments derived from our annotation against those generated by the unsupervised clustering algorithm Clustree at both Day 1 and Day 3 post-fusion. Consistent with our expectations of dynamic changes, we observed significant alterations in both predicted intercellular and intracellular communication networks as the fusion process unfolded over time. Intriguingly, despite the dynamic nature of these networks, the top five ECM-receptor signaling pathways identified in the total population of 38 cells analyzed at Day 3 exhibited remarkable consistency between our manually annotated four cell types and the three cell types independently identified by Clustree. This suggests that while the overall communication landscape undergoes significant remodeling post-fusion, the primary intercellular communication axes of the ECM-receptor signaling become relatively stable by Day 3. The main distinction observed was the order of the Collagen and Thrombospondin pathways within this conserved set of five (**Figure 5B** and **5E**), indicating potential shifts in the relative importance of these specific ECM-mediated interactions.

Our previous study identified an enrichment of cancer-related gene groups in ten transcriptionally distinct hybrid cells compared to their parental mMSC and mHL1 cells. To further explore their potential to exhibit cancerous traits, we had performed PCA using murine breast cancer cell data (MMTV-Wnt-1 TIC-PC and NTC-PC, previously reported^46^), demonstrating that two specific hybrids clustered closely with tumor cell populations, displaying decreased *p53* and elevated *Fos* and *Jun* expression (confirmed by qRT-PCR), even exceeding levels in the breast cancer cells. In the current study, our comparative pathway and TF enrichment analyses of annotated cell types and unsupervised clusters provided further compelling evidence for the acquisition of cancerous characteristics following fusion. Comparative analysis of cell-cell communication pathways revealed dynamic changes between Day 1 and Day 3 of cell culture, with less emphasis on the forementioned breast cancer-related signaling. On Day 1, Wnt signaling and Melanogenesis exhibited reduced enrichment, while Focal adhesion and Endocrine resistance were upregulated in annotation-defined cell types, and Renal cell carcinoma pathways were prominent in Clustree-identified clusters. By Day 3, Endocrine resistance remained elevated in annotation-defined cells, while Wnt signaling showed a more context-dependent influence. Notably, Clustree-identified cells on Day 3 displayed upregulated Focal adhesion and EGFR tyrosine kinase inhibitor resistance, alongside mixed enrichment of Wnt, Estrogen, and Prostate cancer signaling, indicating a shift towards communication involving physical interactions and the emergence of other cancer-associated pathways rather than the abovementioned breast cancer as culture time increased.

While the phenomenon of cellular fusion holds significant promise for potentially reprogramming somatic cells towards an early mesodermal state or generating novel cell types with unique properties, inherent limitations associated with current scRNA-seq technologies necessitate a cautious and nuanced interpretation of our findings. The typical mRNA capture efficiency of 5-10% of the total cellular transcriptome introduces a potential for undersampling, particularly affecting the detection of low-abundance but functionally critical transcripts^47^, such as key TFs involved in lineage commitment and differentiation. This issue is further compounded in the context of hybrid cells, where obtaining true biological replicates can be technically challenging, limiting the statistical power to confidently identify subtle but important gene expression changes. Moreover, our current sample sizes, while providing valuable initial insights, fall below the generally recommended threshold of 500-1,000 cells per sample for robust and statistically powered scRNA-seq analysis, potentially limiting our ability to fully capture the heterogeneity within the fused cell populations and comprehensively model intercellular and intracellular communication networks. To address these limitations and enhance the reliability of our analyses, future research efforts will be strategically directed towards increasing the number of captured single cells per sample. This increase in sequencing depth and cell numbers will enable a more comprehensive and statistically robust investigation of the dynamic changes in gene expression and a more reliable reconstruction of the intricate intercellular and intracellular communication landscapes that govern the behavior and fate of these fusion-derived cell types.

## Conclusions

In conclusion, our investigation into the fusion of mHL1 and mMSC cells has unveiled a rapid and complex rewiring of cellular identity, moving beyond a simple amalgamation of parental traits to generate novel hybrid phenotypes characterized by biased gene expression and dynamic communication networks. Pseudotime analysis highlighted a more protracted transcriptional remodeling in mMSC fusion products, suggesting distinct developmental trajectories post-fusion. Furthermore, the evolution of cell-cell communication pathways revealed a shift over time, with an initial downregulation of Wnt signaling and Melanogenesis giving way to an upregulation of pathways like Endocrine resistance and Focal adhesion, and the emergence of other cancer-associated signaling mechanisms, distinct from the breast cancer pathways, by Day 3. While acknowledging the inherent limitations of current scRNA-seq technologies, our findings underscore the remarkable plasticity of cellular identity following fusion and provide a foundation for future studies aimed at dissecting the precise molecular mechanisms driving these transformations and exploring the potential of cell fusion for generating novel cell types with desired characteristics.

## CRediT authorship contribution statement

**Fateme Nazaryabrbekoh:** Data Curation, Formal Analysis, Investigation, Methodology, Visualization, Writing – Review & Editing.

**JoAnne Huang:** Data Curation, Formal Analysis, Investigation, Methodology, Validation, Visualization, Writing – Review & Editing.

**Syeda S. Shoaib:** Investigation, Methodology.

**Xun Tang:** Investigation, Methodology, Validation.

**Joohyun Kim:** Conceptualization, Methodology, Validation, Writing – Review & Editing.

**Brenda M. Ogle:** Validation, Writing – Review & Editing.

**Jangwook P. Jung:** Conceptualization, Data Curation, Formal Analysis, Funding Acquisition, Investigation, Methodology, Project Administration, Resources, Supervision, Validation, Visualization, Writing – Original Draft Preparation, Writing – Review & Editing.

## Acknowledgement

This work benefited from the support of the National Science Foundation CAREER award DMR 2047018 (JPJ). The authors gratefully acknowledge Brian Freeman for his helpful discussions that contributed to the completion of this study.

