## Supplementary Data for "Cell-Cell Communication and Gene Regulation of Stem Cell-Parenchymal Cell Fusion"

### Supplementary Figures

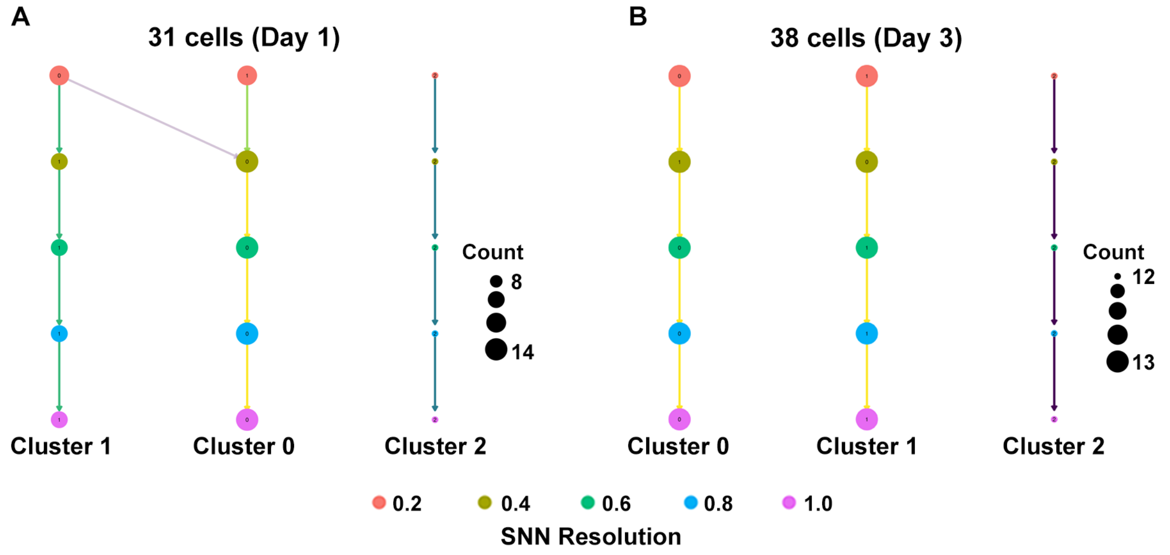


**Figure S1.** Unsupervised clustering of (A) 31 and (B) 38 co-cultured cells is visualized using Clustree. The resulting dendrogram-like structure illustrates the relationships between clusters detected at different SNN resolutions. For each resolution, the size of the points in the rightmost column reflects the number of cells within that designated cluster. Vertically, the figure shows how coarser clusters identified at lower SNN resolutions are progressively refined into more distinct clusters at higher resolutions.

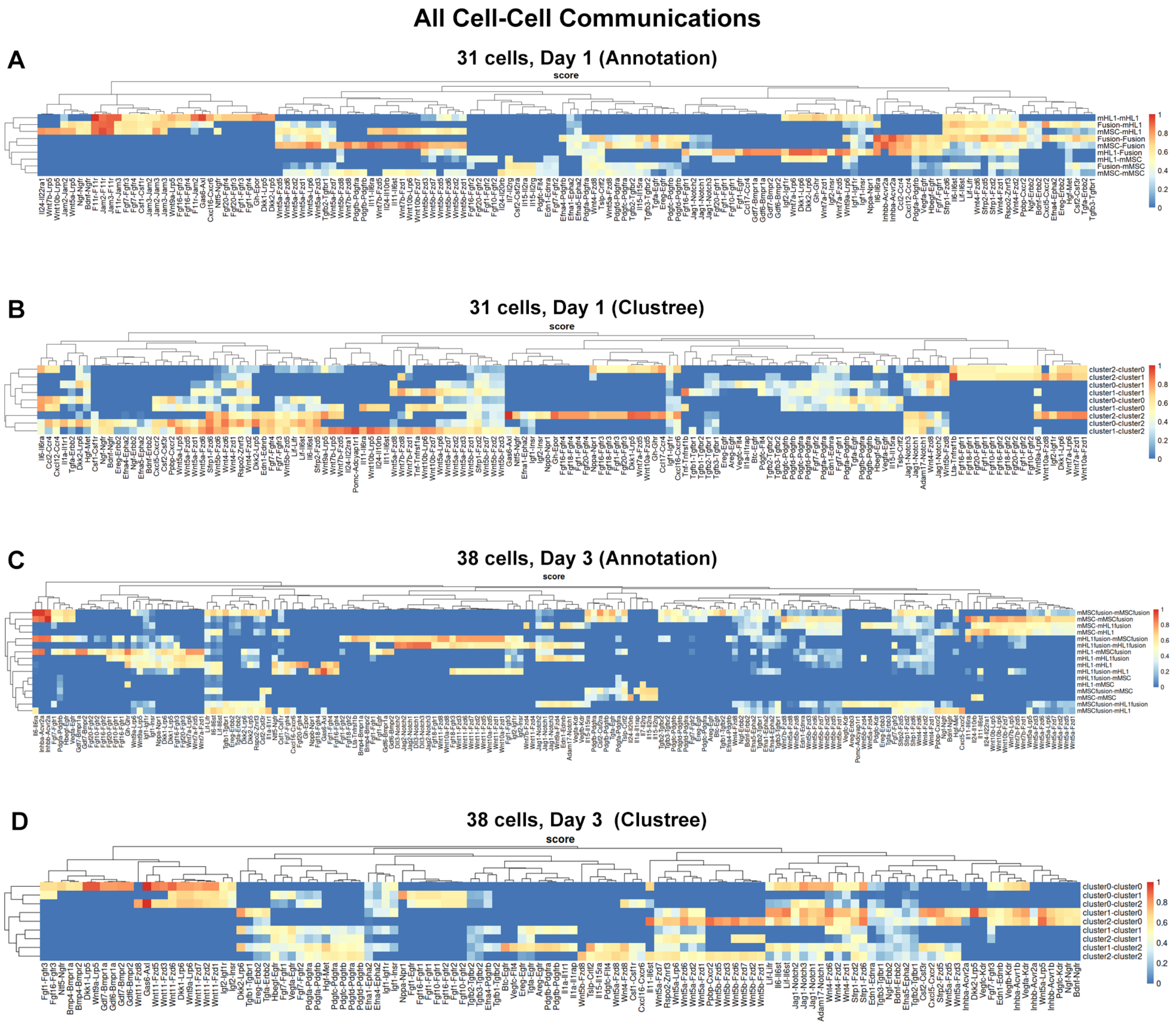


**Figure S2.** Inferred cell-cell communication networks in co-cultured cells at Day 1 (A and B) and Day 3 (C and D) are depicted as heatmaps. Communication scores between cell populations defined by prior annotation are illustrated in (A) and (C), while (B) and (D) present the same data with cell populations organized by unsupervised clustering using Clustree. The color intensity in each heatmap indicates the predicted strength of communication between specific LR pairs or signaling pathways. HC of sending and receiving cell populations is visualized by dendrograms along the top and left of each heatmap. The number of cells analyzed at each time point is specified in the panel titles.

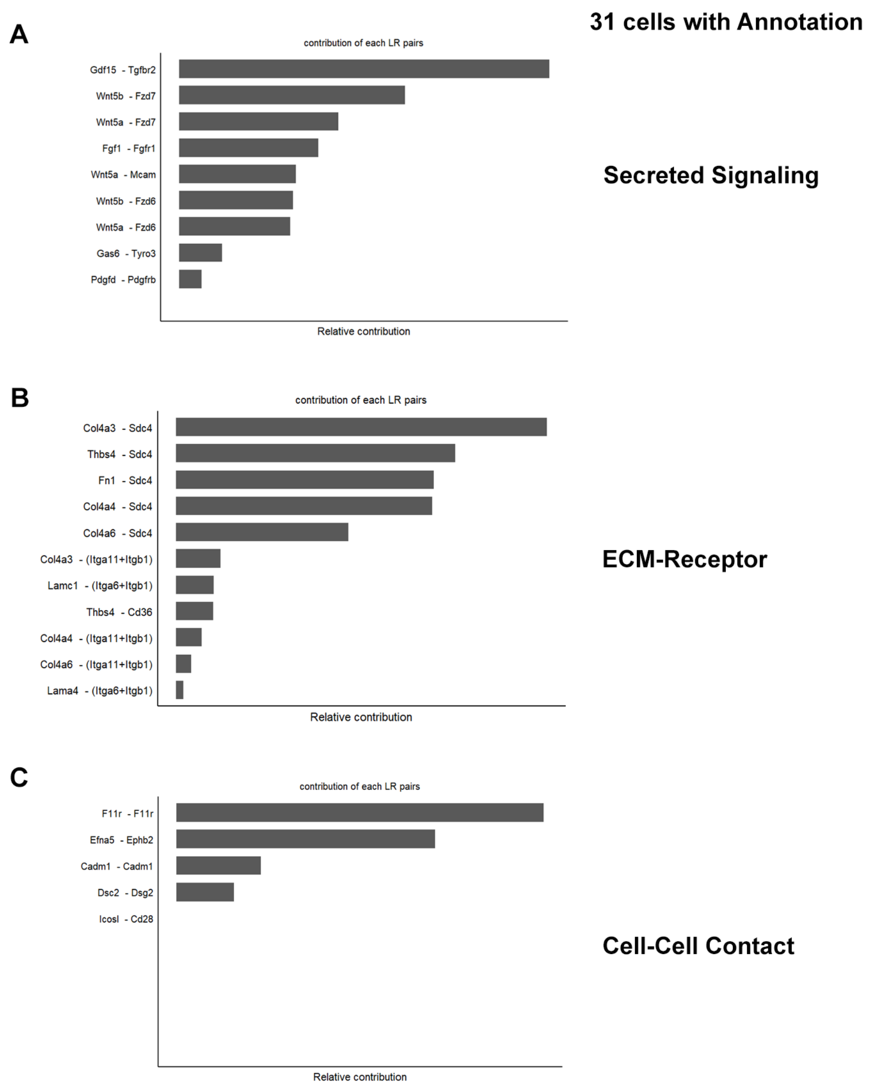


**Figure S3.** Bar plots depict the relative contributions of specific LR pairs to cell-cell communication in 31 annotated co-cultured cells, categorized by communication mechanism. The top contributing LR pairs within Secreted Signaling, ECM-Receptor interactions, and Cell-Cell Contact are detailed in (A), (B) and (C), respectively, with bar length indicating the relative contribution score of each LR pair.

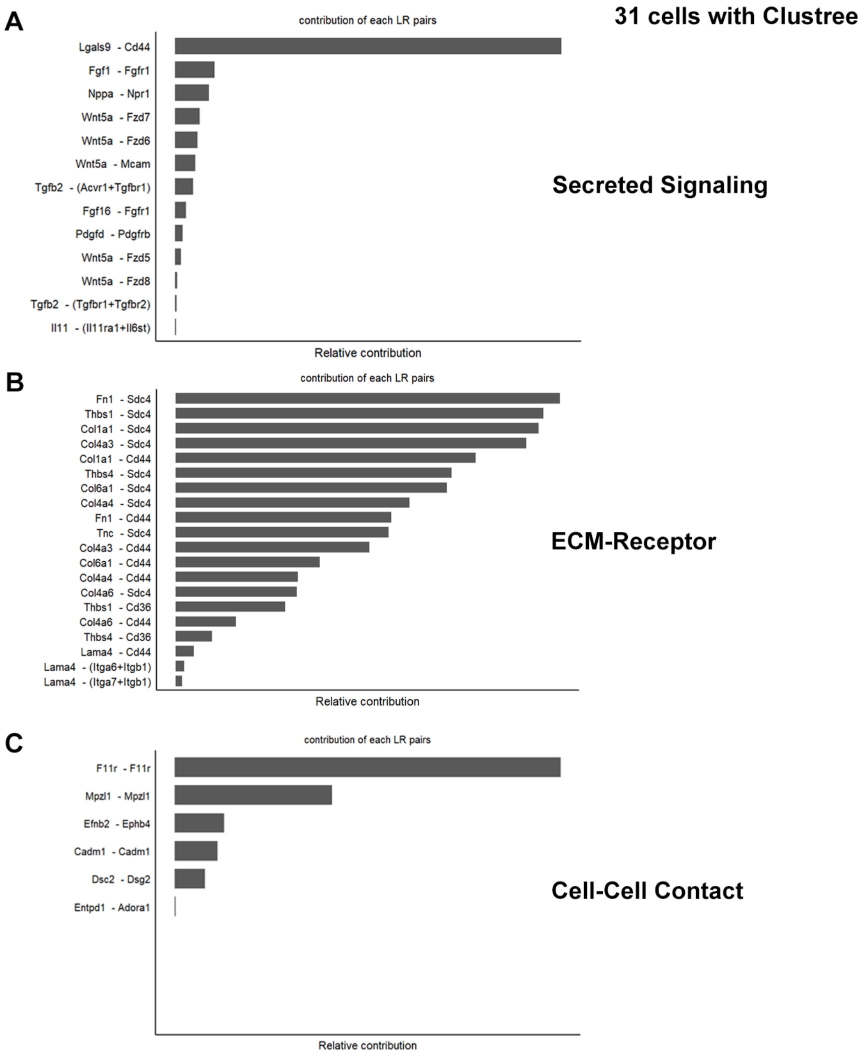


**Figure S4.** Bar plots illustrate the relative contribution of individual LR pairs to cell-cell communication in 31 co-cultured cells, with cell populations defined by unsupervised clustering using Clustree. The top contributing LR pairs within Secreted Signaling, ECM-Receptor interactions, and Cell-Cell Contact are detailed in (A), (B) and (C), respectively.

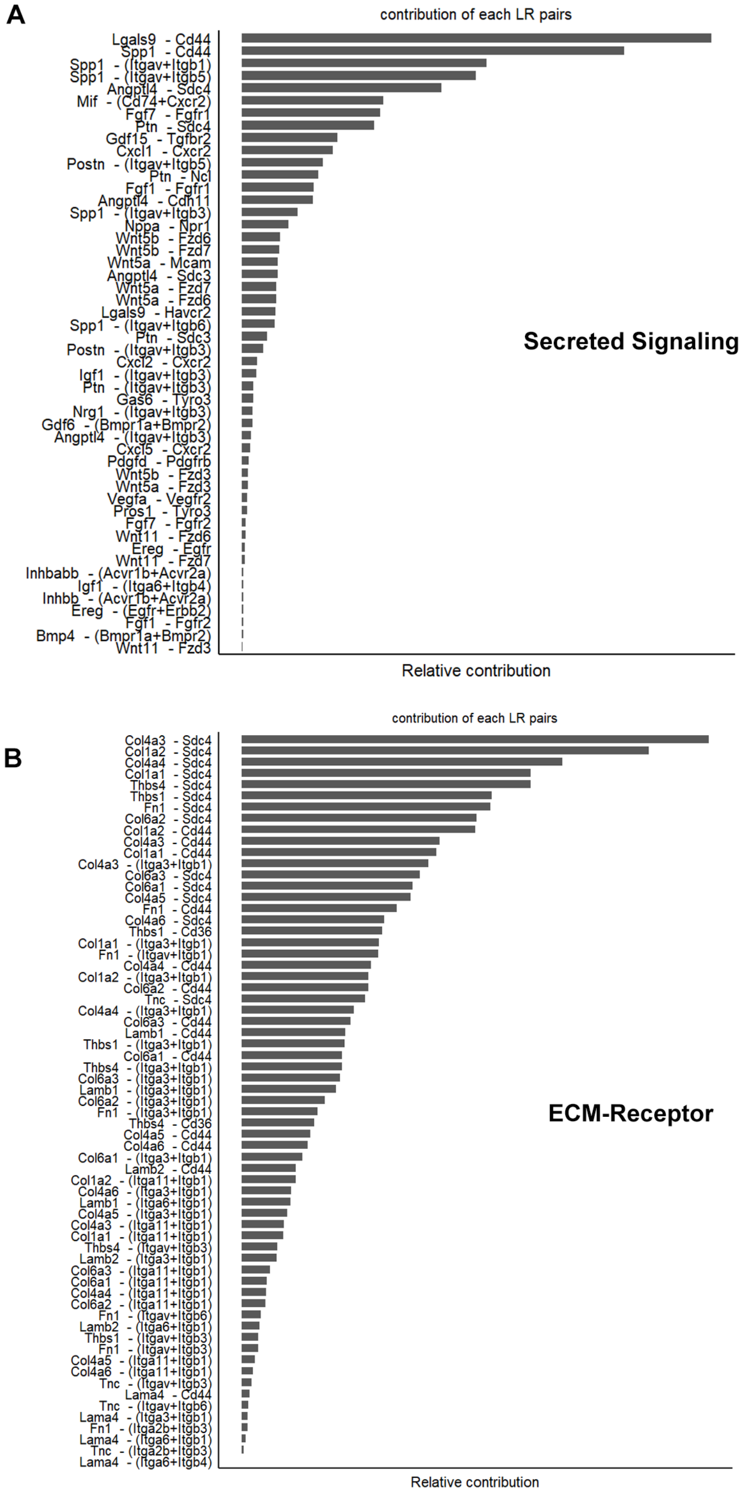
A
contribution of each LR pairs
Lgals9 - Cd44
Spp1 - Cd44
Spp1 - (Itgav+Itgb1)
Spp1 - (Itgav+Itgb5)
Angptl4 - Sdc4
Mif - (Cd74+Cxcr2)
Fgf7 - Fgfr1
Ptn - Sdc4
Gdf15 - Tgfbr2
Cxcl1 - Cxcr2
Postn - (Itgav+Itgb5)
Ptn - Ncl
Fgf1 - Fgfr1
Angptl4 - Cdh11
Spp1 - (Itgav+Itgb3)
Nppa - Npr1
Wnt5b - Fzd6
Wnt5b - Fzd7
Wnt5a - Mcam
Angptl4 - Sdc3
Wnt5a - Fzd7
Wnt5a - Fzd6
Lgals9 - Havcr2
Spp1 - (Itgav+Itgb6)
Ptn - Sdc3
Postn - (Itgav+Itgb3)
Cxcl2 - Cxcr2
Igf1 - (Itgav+Itgb3)
Ptn - (Itgav+Itgb3)
Gas6 - Tyro3
Nrg1 - (Itgav+Itgb3)
Gdf6 - (Bmpr1a+Bmpr2)
Angptl4 - (Itgav+Itgb3)
Cxcl5 - Cxcr2
Pdgfd - Pdgfrb
Wnt5b - Fzd3
Wnt5a - Fzd3
Vegfa - Vegfr2
Pros1 - Tyro3
Fgf7 - Fgfr2
Wnt11 - Fzd6
Ereg - Egfr
Wnt11 - Fzd7
Inhbabb - (Acvr1b+Acvr2a)
Igf1 - (Itga6+Itgb4)
Inhbb - (Acvr1b+Acvr2a)
Ereg - (Egfr+Erbb2)
Fgf1 - Fgfr2
Bmp4 - (Bmpr1a+Bmpr2)
Wnt11 - Fzd3
Secreted Signaling
Relative contribution
B
contribution of each LR pairs
Col4a3 - Sdc4
Col1a2 - Sdc4
Col4a4 - Sdc4
Col1a1 - Sdc4
Thbs4 - Sdc4
Thbs1 - Sdc4
Fn1 - Sdc4
Col6a2 - Sdc4
Col1a2 - Cd44
Col4a3 - Cd44
Col1a1 - Cd44
Col4a3 - (Itga3+Itgb1)
Col6a3 - Sdc4
Col6a1 - Sdc4
Col4a5 - Sdc4
Fn1 - Cd44
Col4a6 - Sdc4
Thbs1 - Cd36
Col1a1 - (Itga3+Itgb1)
Fn1 - (Itgav+Itgb1)
Col4a4 - Cd44
Col1a2 - (Itga3+Itgb1)
Col6a2 - Cd44
Tnc - Sdc4
Col4a4 - (Itga3+Itgb1)
Col6a3 - Cd44
Lamb1 - Cd44
Thbs1 - (Itga3+Itgb1)
Col6a1 - Cd44
Thbs4 - (Itga3+Itgb1)
Col6a3 - (Itga3+Itgb1)
Lamb1 - (Itga3+Itgb1)
Col6a2 - (Itga3+Itgb1)
Fn1 - (Itga3+Itgb1)
Thbs4 - Cd36
Col4a5 - Cd44
Col4a6 - Cd44
Col6a1 - (Itga3+Itgb1)
Lamb2 - Cd44
Col1a2 - (Itga11+Itgb1)
Col4a6 - (Itga3+Itgb1)
Lamb1 - (Itga6+Itgb1)
Col4a5 - (Itga3+Itgb1)
Col4a3 - (Itga11+Itgb1)
Col1a1 - (Itga11+Itgb1)
Thbs4 - (Itgav+Itgb3)
Lamb2 - (Itga3+Itgb1)
Col6a3 - (Itga11+Itgb1)
Col6a1 - (Itga11+Itgb1)
Col4a4 - (Itga11+Itgb1)
Col6a2 - (Itga11+Itgb1)
Fn1 - (Itgav+Itgb6)
Lamb2 - (Itga6+Itgb1)
Thbs1 - (Itgav+Itgb3)
Fn1 - (Itgav+Itgb3)
Col4a5 - (Itga11+Itgb1)
Col4a6 - (Itga11+Itgb1)
Tnc - (Itgav+Itgb3)
Lama4 - Cd44
Tnc - (Itgav+Itgb6)
Lama4 - (Itga3+Itgb1)
Fn1 - (Itga2b+Itgb3)
Lama4 - (Itga6+Itgb1)
Tnc - (Itga2b+Itgb3)
Lama4 - (Itga6+Itgb4)
ECM-Receptor
Relative contribution

C

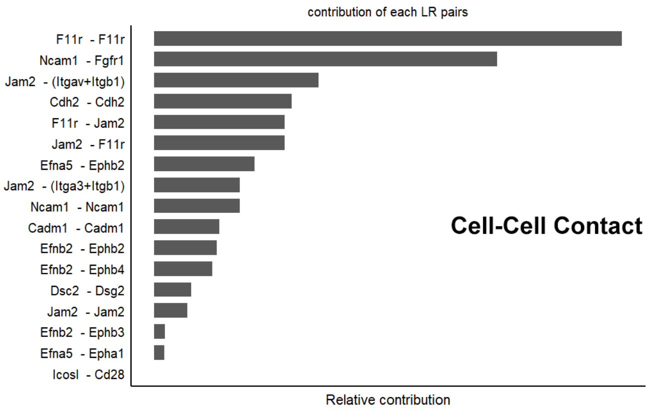


**Figure S5.** Bar plots illustrate the relative contributions of specific LR pairs to cell-cell communication among 38 annotated co-cultured cells, categorized by signaling mechanism. The top contributing LR pairs in Secreted Signaling are shown in (A), and those in ECM-Receptor interactions are highlighted in (B), with bar length representing the relative contribution score within each category. The relative contributions of LR pairs involved in Cell-Cell Contact are specifically detailed in (C).

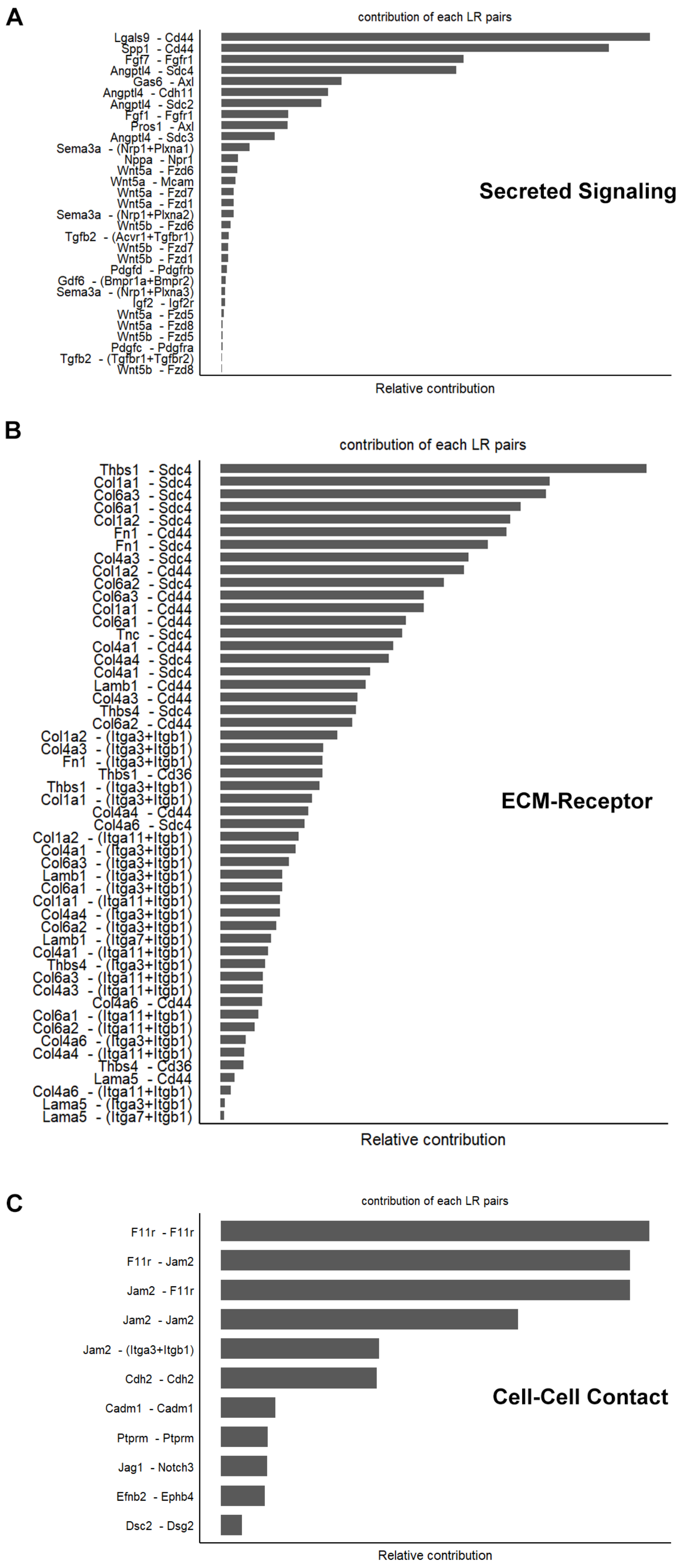
A
contribution of each LR pairs
Lgals9 - Cd44
Spp1 - Cd44
Fgf7 - Fgfr1
Angptl4 - Sdc4
Gas6 - Axl
Angptl4 - Cdh11
Angptl4 - Sdc2
Fgf1 - Fgfr1
Pros1 - Axl
Angptl4 - Sdc3
Sema3a - (Nrp1+Plxna1)
Nppa - Npr1
Wnt5a - Fzd6
Wnt5a - Mcam
Wnt5a - Fzd7
Wnt5a - Fzd1
Sema3a - (Nrp1+Plxna2)
Wnt5b - Fzd6
Tgfb2 - (Acvr1+Tgfbr1)
Wnt5b - Fzd7
Wnt5b - Fzd1
Pdgfd - Pdgfrb
Gdf6 - (Bmpr1a+Bmpr2)
Sema3a - (Nrp1+Plxna3)
Igf2 - Igf2r
Wnt5a - Fzd5
Wnt5a - Fzd8
Wnt5b - Fzd5
Pdgfc - Pdgfra
Tgfb2 - (Tgfbr1+Tgfbr2)
Wnt5b - Fzd8
Secreted Signaling
Relative contribution
B
contribution of each LR pairs
Thbs1 - Sdc4
Col1a1 - Sdc4
Col6a3 - Sdc4
Col6a1 - Sdc4
Col1a2 - Sdc4
Fn1 - Cd44
Fn1 - Sdc4
Col4a3 - Sdc4
Col1a2 - Cd44
Col6a2 - Sdc4
Col6a3 - Cd44
Col1a1 - Cd44
Col6a1 - Cd44
Tnc - Sdc4
Col4a1 - Cd44
Col4a4 - Sdc4
Col4a1 - Sdc4
Lamb1 - Cd44
Col4a3 - Cd44
Thbs4 - Sdc4
Col6a2 - Cd44
Col1a2 - (Itga3+Itgb1)
Col4a3 - (Itga3+Itgb1)
Fn1 - (Itga3+Itgb1)
Thbs1 - Cd36
Thbs1 - (Itga3+Itgb1)
Col1a1 - (Itga3+Itgb1)
Col4a4 - Cd44
Col4a6 - Sdc4
Col1a2 - (Itga11+Itgb1)
Col4a1 - (Itga3+Itgb1)
Col6a3 - (Itga3+Itgb1)
Lamb1 - (Itga3+Itgb1)
Col6a1 - (Itga3+Itgb1)
Col1a1 - (Itga11+Itgb1)
Col4a4 - (Itga3+Itgb1)
Col6a2 - (Itga3+Itgb1)
Lamb1 - (Itga7+Itgb1)
Col4a1 - (Itga11+Itgb1)
Thbs4 - (Itga3+Itgb1)
Col6a3 - (Itga11+Itgb1)
Col4a3 - (Itga11+Itgb1)
Col4a6 - Cd44
Col6a1 - (Itga11+Itgb1)
Col6a2 - (Itga11+Itgb1)
Col4a6 - (Itga3+Itgb1)
Col4a4 - (Itga11+Itgb1)
Thbs4 - Cd36
Lama5 - Cd44
Col4a6 - (Itga11+Itgb1)
Lama5 - (Itga3+Itgb1)
Lama5 - (Itga7+Itgb1)
ECM-Receptor
Relative contribution
C
contribution of each LR pairs
F11r - F11r
F11r - Jam2
Jam2 - F11r
Jam2 - Jam2
Jam2 - (Itga3+Itgb1)
Cdh2 - Cdh2
Cadm1 - Cadm1
Ptprm - Ptprm
Jag1 - Notch3
Efnb2 - Ephb4
Dsc2 - Dsg2
Cell-Cell Contact
Relative contribution

**Figure S6.** Bar plots illustrate the relative contributions of specific LR pairs to cell-cell communication among 38 cells, with cell populations defined by Clustree. Secreted Signaling is depicted in (A), and ECM-Receptor interactions are shown in (B); in both, bar length corresponds to the relative contribution score of each LR pair to the overall communication within that category. The relative contributions of LR pairs involved in Cell-Cell Contact are specifically illustrated in (C).

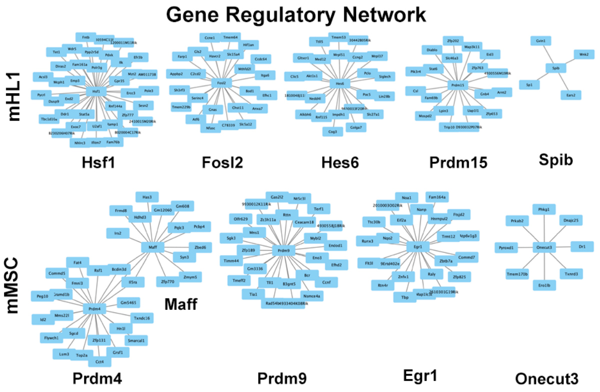


**Figure S7.** GRNs for selected TFs in mHL1 and mMSC cell types. Each sub-network depicts the regulatory relationships between a central TF (labeled below each network) and its predicted target genes (connected nodes), with arrows indicating the direction of regulation (TF regulating target gene).

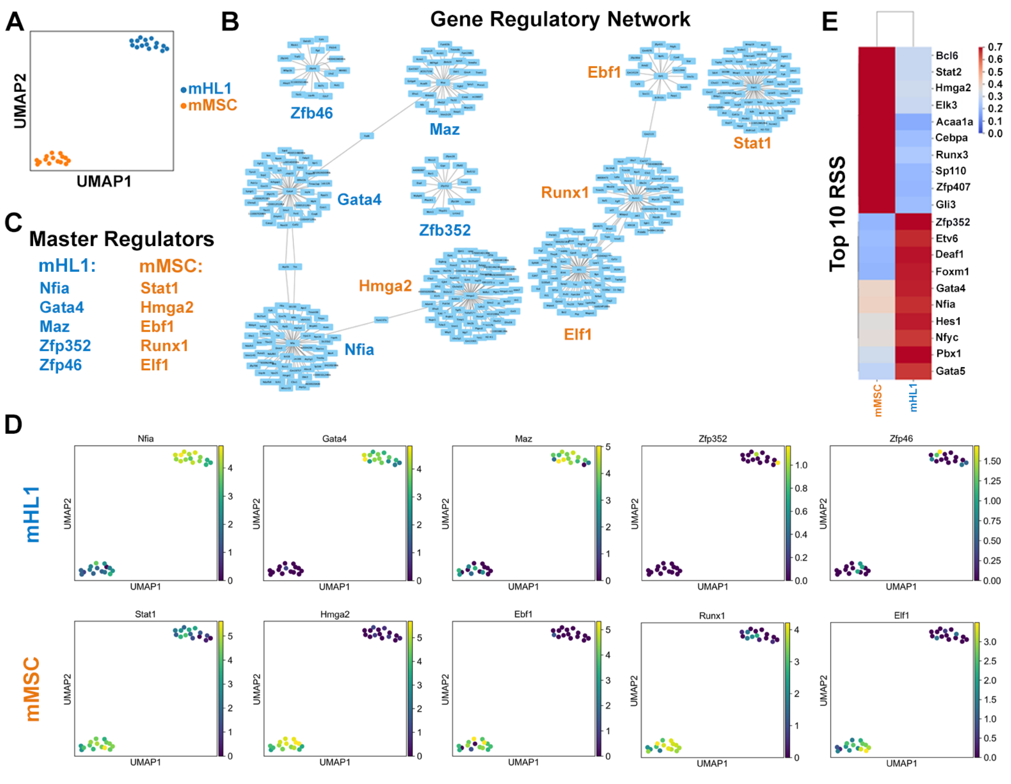


**Figure S8.** GRN analysis of mHL1 and mMSC cell types. (A) UMAP visualization showing the distribution of mHL1 (blue) and mMSC (orange) cells. (B) Network of master regulator TFs identified in mHL1 and mMSC, with edges representing potential regulatory relationships. (C) List of the top master regulator TFs identified for each cell type. (D) Feature plots on the UMAP embedding showing the expression levels of selected master regulator TFs in mHL1 and mMSC cells, with color intensity indicating expression level. (E) Heatmap displaying the RSS of the top 10 master regulator TFs for each cell type, indicating the degree to which their regulon activity is specific to either mHL1 or mMSC.

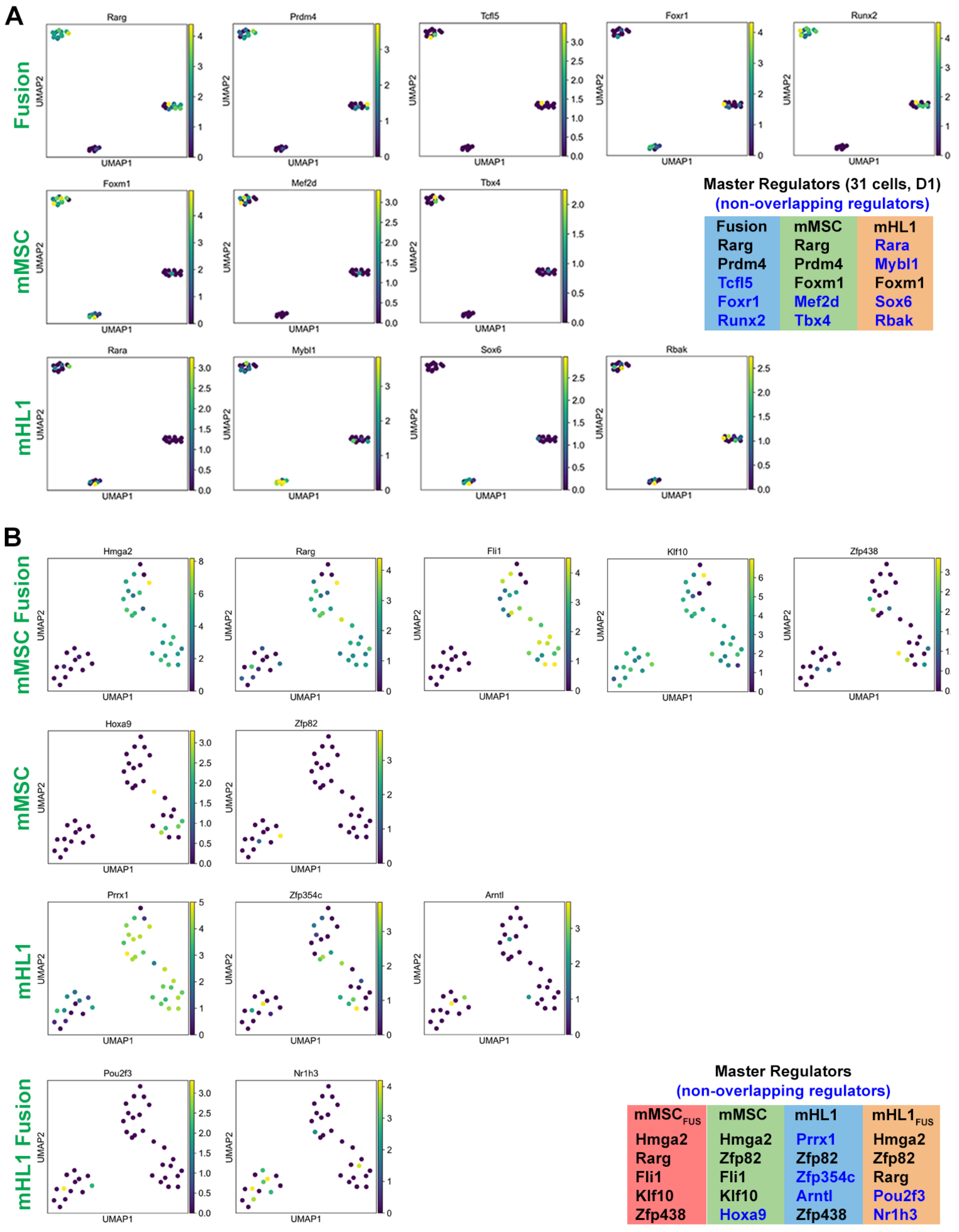


**Figure S9.** UMAP visualizations displaying the expression patterns of selected master regulator TFs in Fusion, mMSC and mHL1 cell populations (31 cells, Day 1). (A) Feature plots showing the expression levels of TFs identified as master regulators in Fusion, mMSC and mHL1 cells. Color intensity indicates the expression level of each TF within the UMAP embedding. (B) Feature plots illustrating the expression levels of additional master regulator TFs identified in the respective cell populations. The legend provides a key to the TFs specific to each cell type.

### Supplementary Tables

#### Table S1. Tabula Muris cell types per module.

| | |
|---|---|
| **Partition mMSC D0** | |
| Module 30 | Granulocyte, Fat CL:0000094 |
| Module 27 | Natural Killer Cell, Marrow CL:0000623 |
| Module 12 | Fibroblast, Heart CL:0000057 |
| Module 13 | Mesenchymal cell, bladder CL:00008019 |
| Module 11 | Mesenchymal cell, bladder CL:00008019 |
| **Partition D1** | |
| Module 8 | Endothelial cell, kidney CL:0000115 |
| Module 17 | Fraction A Pre Pro B Cell, Marrow CL:0002045 |
| Module 26 | Mesenchymal cell, bladder CL:00008019 |
| Module 1 | Mesenchymal stem cell, muscle CL:0000134 |
| Module 19 | Epithelial cell, large intestine Colon CL:0002253 |
| **Partition mMSC D1** | |
| Module 15 | Stromal Cell, Trachea CL:0000499 |
| Module 17 | Fraction A Pre Pro B Cell, Marrow CL:0002045 |
| Module 10 | Basal Cell of Epidermis, Tongue CL:0002187 |
| Module 19 | Epithelial cell, large intestine Colon CL:0002253 |
| Module 6 | Skeletal Muscle Satellite Cell, Muscle CL:0000594 |
| **Partition mHL-1 D0** | |
| Module 9 | Stem Cell of Epidermis, Skin CL:1000428 |
| Module 24 | Cardiac Muscle Cell, Heart CL:0000746 |
| Module 5 | Cardiac Muscle Cell, Heart CL:0000746 |
| Module 7 | Stem Cell of Epidermis, Skin CL:1000428 |
| Module 28 | Stem Cell of Epidermis, Skin CL:1000428 |
| **Partition mHL-1 D1** | |
| Module 29 | Keratinocyte, Tongue CL:0000312 |
| Module 18 | Stromal Cell, Trachea CL:0000499 |
| Module 2 | Stem Cell of Epidermis, Skin CL:1000428 |
| Module 4 | Stem Cell of Epidermis, Skin CL:1000428 |
| Module 3 | Cardiac Muscle Cell, Heart CL:0000746 |
| **Partition D3** | |
| Module 29 | Keratinocyte, Tongue CL:0000312 |
| Module 21 | Smooth muscle cell, fat CL:0000192 |
| Module 23 | Skeletal Muscle Satellite Cell, Muscle CL:0000594 |
| Module 22 | Cardiac Muscle Cell, Heart CL:0000746 |
| Module 3 | Cardiac Muscle Cell, Heart CL:0000746 |
